# HIV infection in microglia leads to senescence, triggering activation of neurotoxicity pathways

**DOI:** 10.1101/2025.05.08.651477

**Authors:** Sara J. Mason, Sheetal Sreeram, Farshad Niazi, Konstantin Leskov, Alan D. Levine, Jonathan Karn, Saba Valadkhan

## Abstract

HIV-associated neurocognitive disorders (HAND) persist in milder forms despite anti-retroviral therapy, leading to premature and exacerbated aging-related cognitive disorders. We investigated the interplay between HAND and aging in microglia, which constitute the main brain HIV reservoir. We compared the transcriptomic patterns associated with normal aging in healthy humans to those observed following HIV infection in both ex vivo and in vivo models. Single cell and bulk transcriptomic patterns revealed that HIV infection induces a pattern of cellular senescence, with strong parallels to the transcriptomic signature of normal aging. Both processes were characterized by p53 pathway activation, upregulation of inflammatory genes and downregulation of proliferative genes while maintaining mTOR signaling, a pattern characteristic of cellular senescence. Importantly, both actively HIV infected and bystander microglia showed the cellular senescence patterns. Our results provide a mechanistic basis for the observed premature brain aging in HAND, and identify senescence-associated pathways as potential therapeutic targets.

## Introduction

Despite decades of scientific advancement and the development of highly effective combination antiretroviral therapy (cART), HIV represents a prime example of persistent viral infection that continues to challenge our understanding of virus-host interactions, immune evasion, and long-term pathogenesis across multiple organ systems. While cART can effectively control viremia in people living with HIV (PLWH), HIV-associated impairment of sensory, motor, and cognitive processes in the brain, collectively known as HIV-associated Neurocognitive Disorders (HAND) remain a major health challenge^1–14^. Although the prevalence of the most severe form of HAND, HIV-associated Dementia (HAD) has strongly decreased with the use of cART and is observed in 2-5% of PLWH, milder forms of HAND including Asymptomatic Neurocognitive Impairment (ANI) and Mild Neurocognitive Disorder (MND) are observed in up to 26% and 15% of PLWH, respectively^15,16^. In adults over the age of fifty, PLWH had over seven fold increased prevalence in mild cognitive impairment compared to non-PLWH individuals ^17^.

While long-term cART exposure may have some neurotoxic effects, HIV-associated brain injury is thought to play the dominant role in the development of HAND^18^. Mechanistically, HIV-associated brain injury results from a multifaceted interplay of direct viral protein neurotoxicity and chronic neuroinflammation, leading to blood-brain barrier dysfunction, impaired neurogenesis, oxidative stress, excitotoxicity, synaptic dysfunction, and subsequent neuronal damage^19–22^. HIV-infected microglia, which serve as the primary HIV reservoir in the brain, play a key role in the pathogenesis of HAND and the induction of HIV-associated brain injury^23^. Direct HIV infection of microglia leads to chronic activation and the release of inflammatory mediators, which in turn induce the majority of the pathological changes listed above. Further, microglia serve as long-lived reservoirs for HIV in the brain, contributing to ongoing viral persistence despite anti-retroviral therapy (ART)^24^.

While HIV crosses the blood-brain barrier and infects microglia early in the course of infection, the development of HAND is not solely dependent on the presence of the infected microglia in the brain. Rather, the variability in HAND prevalence depends on additional factors modulating the risk and severity of neurocognitive impairment^25^. One of the strongest predictors of the development of HAND is age, with the prevalence of HAND in PLWH aged 50 years and older reaching as high as 47%, along with a threefold increased risk of HIV-associated dementia (HAD) compared to younger PLWH^1,15,25–33^. It is widely thought that age and HIV synergistically increase the risk of neurocognitive decline and may interact to lower cognitive reserve^21^, at least partly through inducing premature or accelerated brain aging^20,28,34–36^. Evidence for HIV-mediated accelerated aging range from the presence of epigenetic changes in human brain and blood samples from PLWH^37^, altered IgG glycan profiles^38^, and at the cellular level, increased senescence-associated β-galactosidase activity, elevated p21/CDKN1A and p16/CDKN2A expression in ex-vivo HIV-infected human microglia^36–40^. Together, the available evidence strongly suggest that HIV infection likely leads to a senescence-like phenotype in microglia.

At the cellular level, senescence is defined as sustained elevation of the p53 pathway in the absence of reduced mTOR signaling and is associated with elevated expression of p53-p21 axis and/or p15/p16 pathway leading to stabilized cell cycle arrest; permanent DNA damage foci, and increased activity of the senescence-associated β-galactosidase^36,41–45^. Cellular senescence is also associated with the senescence-associated secretory phenotype (SASP), which involves the release of pro-inflammatory cytokines and metalloproteinases that are implicated in neuroinflammation and neurotoxicity^46–49^. The release of inflammatory factors and extracellular matrix-modifying enzymes can affect neighboring cells, potentially propagating inflammation and cellular dysfunction beyond the originally senescent cells^48,49^. Given these implications, it is crucial to firmly establish whether HIV infection induces a bona fide senescence state in microglia, as it would significantly impact our understanding of HAND pathogenesis and potentially reveal novel therapeutic targets for intervention.

To define the global impact of HIV infection on microglia, we performed an in-depth transcriptomic study on bulk and two single-cell RNA-seq (scRNA-seq) datasets from HIV-infected and mock infected human induced pluripotent stem cells (iPSC) derived microglia on days 7, 15 and 29 post-infection^50^. We found that HIV infection in microglia led to activation of multiple inflammatory pathways including TNF-NFkB, IL6-JAK-STAT and IL1B signaling along with the interferon (IFN) response. Activation of the p53 pathway and the induction of multiple cellular senescence markers resulted in global downregulation of proliferative pathways and the associated regulatory networks, along with the downregulation of antisenescence networks such as Sirtuin1 and NORAD^51–56^. We used transcriptomic databases of primary human microglia including both younger and older individuals to identify a transcriptomic signature for microglial aging. Importantly, our analyses showed this microglial aging signature pattern to be prominently present in HIV-infected cells. These results, in aggregate, prove that HIV infection leads to cellular senescence in microglia. Analysis of two scRNA-seq datasets from iPSC-derived, HIV infected microglial models indicated that cells showing the expression of HIV genes had the strongest signature of senescence and expression of inflammatory cytokines. However, cells exposed to HIV but with low or no detectable HIV gene expression, corresponding to latently infected or uninfected bystander cells, also showed a clear senescent phenotype, suggesting that the bystander cells also contribute to the pathogenesis of HAND. To determine whether the HIV-induced senescence is also observed in vivo, we used scRNA-seq from SIV-infected macaque models^57^. Not only the SIV-infected and bystander microglia showed a clear pattern of senescence, but also residual senescence patterns were present in microglia from SIV-infected animals who received a clinically relevant cART regimen. This surprising result indicates that despite the absence of the main hallmarks of senescence in cells from treated animals, cART can’t reverse all pathogenic impacts of infection. Importantly, data from trajectory analysis and in vivo surface interactome studies indicate that senescence is associated with loss of neuroprotective gene expression and induction of neurotoxic signaling, thus providing a senescence-mediated mechanism for the development of HAND. These findings not only provide a novel mechanistic explanation for the observed inflammatory activation of microglia and exacerbation of brain aging in PLWH, but also point to senescence-related mechanisms as a promising avenue for identification of novel therapeutic and preventative strategies for HAND.

## Results

### HIV infection leads to cellular senescence in human iPSC-derived microglia

In order to understand the global transcriptomic changes induced in microglia following HIV infection, we took a transcriptomic approach using a publicly available RNA-seq dataset from iPSC-derived microglia (iMg) from three donors^50^. iMG cells were infected with JAGO strain of HIV or mock-infected, followed by maintenance in monoculture until harvested for bulk RNA-seq on day 15 post-infection, as described^50^. We first confirmed the purity of the microglial population using microglial marker genes, including P2RY12, CX3CR1, and AIF1, as well as the absence of expression of astrocyte, neuron, and oligodendrocyte markers (Supplemental Figure 1A). HIV infected populations showed a robust expression of HIV genes (supplemental figure 1B), while largely maintaining the expression of microglial markers.

To identify changes in major cellular pathways in microglia in response to HIV infection, we performed pathway analysis on over 1300 genes changing in expression by at least two-fold (FDR<0.05) in response to HIV infection which we had identified by pairwise differential gene expression analysis (Supplemental figure 1C). The most prominently enriched pathways were inflammatory pathways, including the TNFα-NFkB axis, IL6-JAK-STAT and IFN pathways (Figure 1A). The enrichment of these pathways were driven by TNF- and IFN-induced genes in addition to IL1B, a key driver of inflammation in microglia, together with multiple cytokines including CCL3, CCL3L1 and CXCL16 among several other genes (Figure 1B). The p53 pathway was the second most positively enriched, accompanied by negative enrichment of multiple cell-cycle related pathways, including the G2M checkpoint, E2F targets, mitotic spindle, and MYC pathways (Figure 1A). Within these pathways we identified a strong decrease in genes involved in positive regulation of cell proliferation including cyclins, cyclin-dependent kinases, and genes involved in DNA replication and mitosis along with upregulation of anti-proliferative genes, which pointed to a sharp halt in cellular proliferation in HIV-infected microglia (Figure 1C, D, E, F, Supplemental Figures 2, 3). Gene regulatory network analysis also showed a strong decrease in the expression of key transcription factors involved in initiating cell cycle progression and their target genes (Figure 1G). To gain insight into the changes in expression of genes forming the core regulatory elements of cellular pathways, we performed pathway analysis using KEGG gene lists. Intriguingly, in addition to confirming the observed enrichment of inflammatory pathways and downregulation of proliferative ones, our analysis pointed to a marked perturbation of the cellular senescence pathway (Figure 2A, Supplemental Fig. 4). Analysis of the expression pattern of genes involved in cellular senescence indicated a strong upregulation of genes involved in inducing cell cycle arrest and downregulation of proliferative genes, along with extensive remodeling of pathways involved in oxidative and mitochondrial stress, inflammatory signaling, tissue damage, metabolic stress, impaired autophagy and apoptosis resistance (Fig. 2B, C, Supplemental Fig. 4). Together, these changes in gene expression pattern were consistent with a strong induction of the classic DNA-damage-driven cellular senescence process induced by the dual activation of the p53 and NFkB axes (Fig. 1A). Cell cycle block initiated by p53, p21/CDKN1A and GADD45 was reinforced and sustained through the upregulation of p15/CDKN2B, which leads to the engagement of the RB axis. On the other hand, NFkB activation induces the inflammatory response and tissue damage process through SASP which involves the enhanced secretion of multiple cytokines and growth factors, remodeling of the extracellular matrix, along with metabolic rewiring, autophagy stress (p62/SQSTM1 upregulation), apoptosis resistance (CASP3 downregulation) and loss of anti-senescence signaling (SIRT1)^58,59^. Importantly, despite the strong activation of p53 pathway and cell cycle arrest, we did not observe a downregulation in mTOR pathway, ruling out the entry of cell cycle arrested cells into quiescence (Supplemental Fig. 5)^38–42^. We also analyzed the expression of long non-coding RNAs (lncRNAs), a highly under-studied class of regulatory molecules known to play critical roles in regulation of cellular function. We observed an increase in PVT1, an lncRNA target of p53, which blocks MYC expression; along with a decrease in NORAD, a lncRNA whose downregulation is known to induce cellular senescence in both humans and mice^51–53,60^. Taken together, these results indicate that HIV infection in microglia results in the strong induction of cellular senescence through multiple axes of regulation (Fig. 2D).

**Figure 1.**
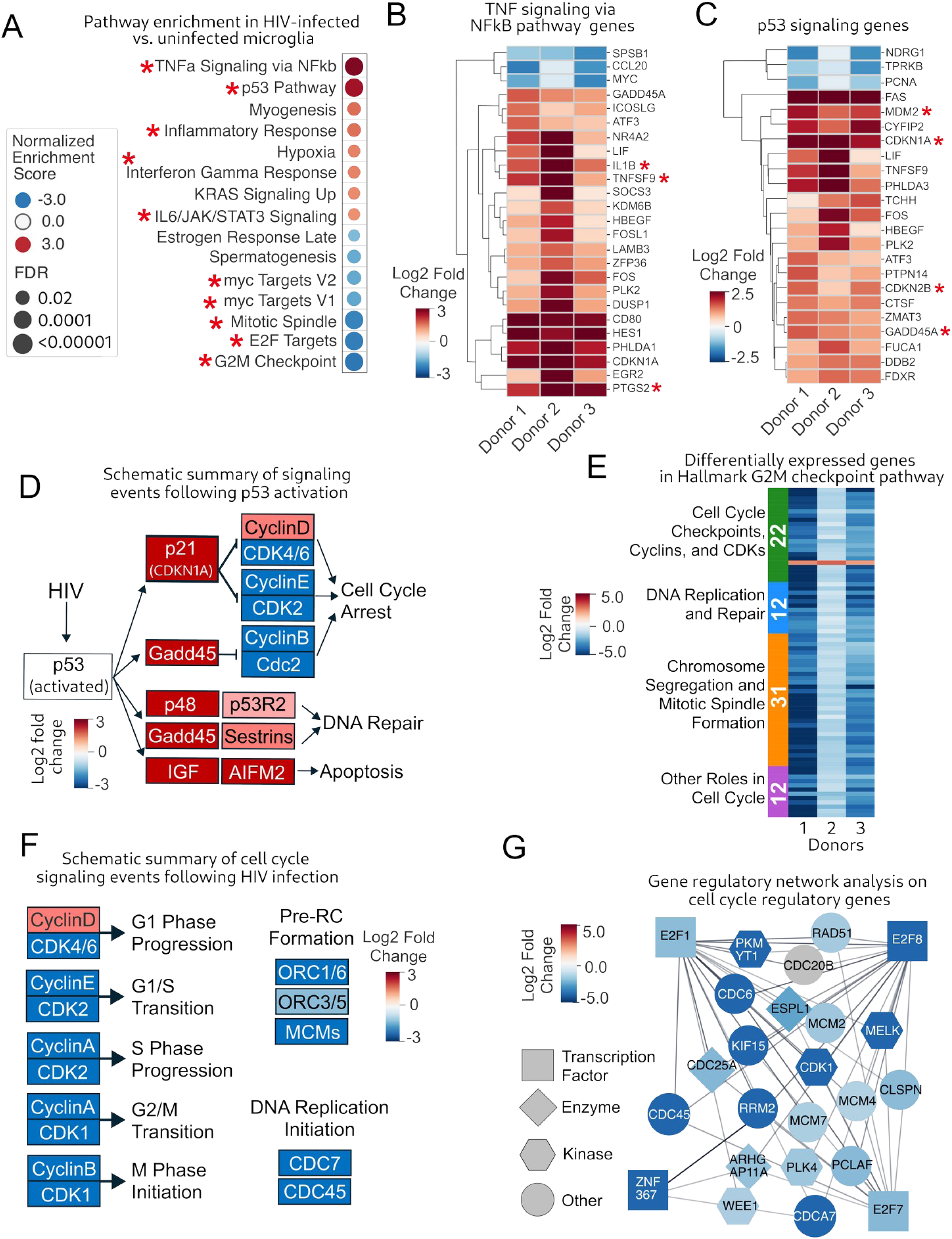
HIV infection leads to the induction of inflammatory pathways and cell cycle arrest. A. Dotplot of hallmark pathways increasing or decreasing in microglia in response to HIV infection. B. Heatmap of genes differentially expressed in microglia in response to HIV infection in the hallmark TNFa signaling via NFkB pathway. Asterisks denote key pro-inflammatory genes. C. Heatmap of genes differentially expressed in microglia in response to HIV infection in the Hallmark p53 signaling pathway. Asterisks denote key targets of p53. D. Diagram of genes changing in microglia in response to HIV in the KEGG p53 pathway. E. Heatmap of genes differentially expressed in microglia in response to HIV infection in the Hallmark G2M checkpoints pathways. F. Diagram of genes changing in microglia in response to HIV infection in the KEGG cell cycle pathway. G. Gene regulatory network analysis including major transcription factors involved in cell cycle regulation and their targets in microglia in response to HIV infection.

**Figure 2.**
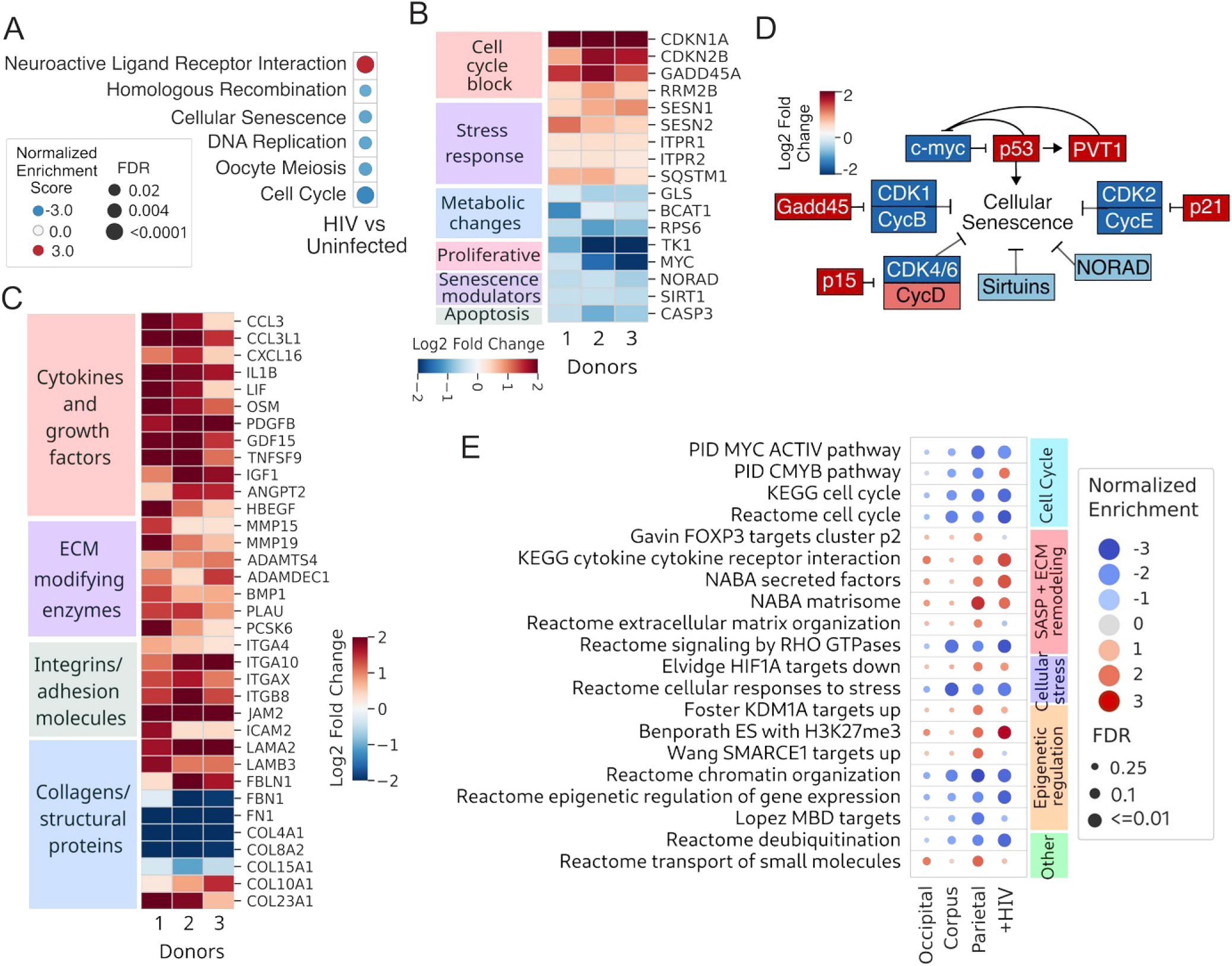
HIV infection induces cellular senescence in microglia. (A) Dotplot of KEGG pathways increasing or decreasing in microglia in response to HIV infection. (B) Diagram of genes changing in response to HIV infection in microglia in the Hallmark mTOR pathway. (C) Heatmap of lncRNAs differentially expressed in microglia in response to HIV infection. (D) Diagram of genes changing in microglia in response to HIV in the KEGG cellular senescence pathway. (E) Heatmap of genes correlating positively or negatively with age in microglia in the occipital cortex and corpus callosum in the Hallmark TNFa siganling via NFkB pathway. (F) Heatmap of genes correlating with age in microglia in the occipital cortex and corpus callosum in the Hallmark G2M checkpoint and E2F targets pathways. (G) Dotplot of core microglial aging pathways in all three brain loci and iPSC-derived microglia infected with HIV. The pathways positively or negatively enriched in HIV-infected microglia were directly matched against the top 20 microglial aging signature pathways (shown in the three columns of dots to the left). 16 of 20 pathways were concordantly enriched in HIV-infected cells.

### HIV infection leads to gene expression patterns mimicking age-associated changes in primary human microglia

The link between natural aging, which leads to the development of cellular senescence, and the development or worsening of HIV-associated neurocognitive disorders is well-established ^1,61,62^. To investigate whether the age-related exacerbation of HAND results from synergistic amplification of shared senescence-associated gene expression patterns and downstream pathogenic processes, we analyzed gene expression profiles associated with natural aging. We used two bulk RNA-seq datasets from post-mortem primary human microglia which were isolated from three brain regions (corpus callosum, occipital and parietal cortex) of healthy donors aged 34-102 years^63,64^. After confirming the purity of the microglial populations (supplemental figure 6A, B), we identified genes positively or negatively correlated with age in each of the three brain regions in our study, namely the parietal cortex, corpus callosum, and occipital cortex (supplemental figure 6C). As microglial phenotypes differ based on the brain region they originate from, the three analyses were compared to define a core microglial signature of aging, consisting of genes and pathways that correlate with age concordantly in at least two of the three brain loci (Fig. 2E, supplemental Fig. 7A, B). We observed that the genes increasing in expression with age were enriched in inflammatory pathways along with extracellular matrix remodeling (potentially representing SASP genes) and epigenetic pathways, known to play crucial roles in the senescence process^65–67^ (Figure 2E). On the other hand, downregulated genes were members of cell cycle-related, metabolic, and senescence-modulating pathways (Figure 2E). To define the similarities between HIV-infected microglia and aged ones, the enrichment pattern of pathways we had identified as part of the core microglial signature of aging (Fig. 2E, the left three columns) was directly assessed in the HIV-infected microglia. Interestingly, of the top 20 aging-enriched pathways, 16 of them showed concordant enrichment in HIV infected cells, indicating close parallels between aged and HIV-infected microglia (Figure 2E). Strong agreement was seen in cell cycle, inflammatory cytokine secretion, extracellular matrix remodeling, stress response and epigenetic processes, pathways which comprise the signature of cellular senescence (Figure 2E). The presence of senescence signature processes in both aged microglia and HIV-infected ones suggests that HIV infection and aging likely converge on cellular senescence pathways, potentially acting synergistically to exacerbate age-associated microglial changes. Given that dysregulated microglial function is an established contributor to the pathogenesis of HAND, these findings provide a molecular basis for understanding how HIV infection might accelerate neurocognitive decline in aging individuals.

### Both actively infected and bystander microglia show signatures of senescence

While our analysis of bulk RNA-seq datasets of iPSC-derived microglia cultures (in which the vast majority of cells were actively infected) showed a clear senescence phenotype at the population level, the percentage of HIV-infected microglia in PLWH is estimated to be much lower, at approximately 1-2%^23^. To determine whether the observed senescence signature is only present in microglia actively infected with HIV, or latently infected and uninfected, bystander microglia also show senescence phenotypes, we utilized single cell RNA-seq (scRNA-seq). We derived microglial cells from human iPSCs, as described by Abud et.al in monocultures. Briefly, the iMGs were differentiated for 27 days and infected with the replication-competent, macrophage R5-tropic, NL-AD8 HIV-1. Uninfected iMGs and HIV-infected iMGs at day 7 post-infection, were used for scRNA-seq. The resulting dataset was analyzed as described^68^. Clustering analysis revealed the presence of three clusters (Fig. 3A), all of which showed a strong expression of microglial lineage markers and a cellular population expressing HIV genes (Fig. 3A, B, Supplementary Fig. 8A). Interestingly, comparison of the gene expression patterns of the three clusters indicated that while clusters 0 and 1 corresponded to homeostatic and inflamed microglia, cluster 2 showed a strong increase in the expression of p16/CDKN2A, a marker of senescence (Fig. 3C, Supplemental Fig. 8B, C). Importantly, the infected sample had a higher proportion of p16/CDKN2A-positive cells (5.91% and 9.20% in mock infected and infected cells, respectively, Fig. 3C). Analysis of the expression of additional senescence markers including p21/CDKN1A and p62/SQSTM1 also showed enriched expression in cluster 2, in addition to the inflamed cluster 1 (Fig. 3D). Further, several SASP genes, including IL8, IL1A, IL1B, CCL2/3/4 also showed enriched expression in clusters 1 and 2 (Fig. 3D).

**Fig. 3.**
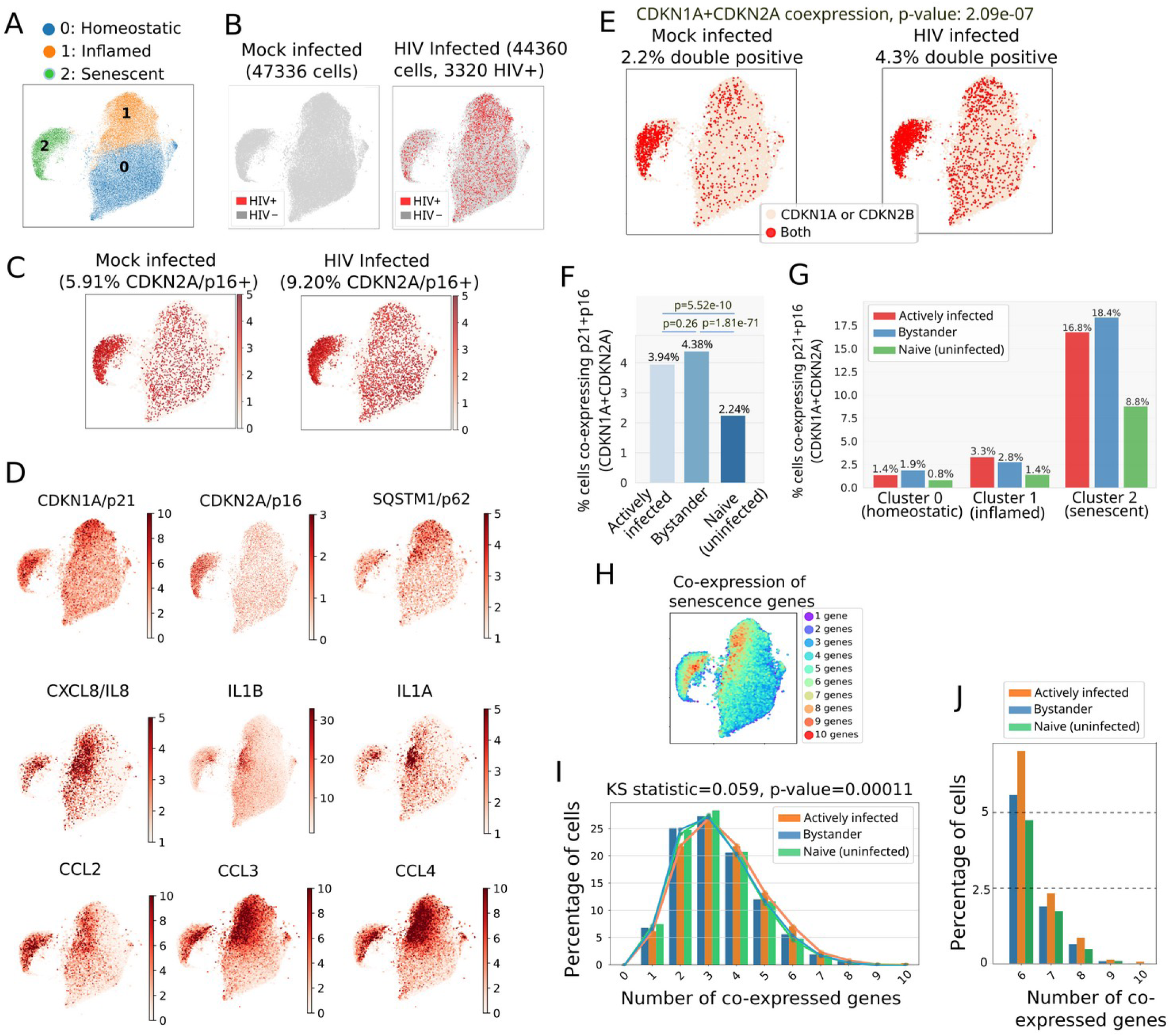
HIV infection induces senescence in both actively infected and bystander cells. A. UMAP of the clusters corresponding to homeostatic, inflamed and senescent cells. B. UMAPs of infected and mock infected cells. Red dots point to HIV+ cells. The number of cells in each group is shown above each panel. C. Compared to the naive, mock-infected sample, the sample exposed to HIV has nearly twice the number of p16/CDKN2A+ cells. D. Markers of senescence (top row) and SASP (lower two rows) are actively expressed in clusters 1 and 2. E. Strong co-expression of p21/CDKN1A and p16/CDKN2B in infected cluster 2 points to the induction of senescence in this cluster. p-value represents the statistical significance of increased p21/p16 co-expression in the HIV-exposed vs mock infected cells. F. Co-expression of p21/p16 is close to two fold higher in cells actively infected and bystander cells. G. Within all three clusters, the percentage of cells co-expressing p21/p16 is close to two fold higher in HIV exposed cells compared to naive, untreated cells. H. Extent of co-expression of the senescence markers and SASP genes. At least one of p21/p16 hallmark senescence markers are present in the cells included in the co-expression analysis. I. Co-expression patterns of senescence markers and SASP genes in naive, bystander and activelyinfected cells shows a shift in the actively infected cells. The Kolmogorov-Smirnov (KS) statistic and p-value refer to comparison of actively infected vs. naive cells. J. Cells with six or more co-expressed genes are enriched in actively infected cells.

In order to directly assess the extent of HIV-induced senescence, we used the commonly accepted framework in which the combined presence of two primary senescence signature genes (i.e. p21/CDKN1A, p16/CDKN2A or p15/CDKN2B), or the presence of at least one of them along with multiple supplementary markers (such as p27/CDKN1B, p62/SQSTM1, in addition to multiple SASP-related genes) is used to define the senescent state^69–72^. As co-expression of CDKN1A and CDKN2A leads to stable induction of cell cycle arrest and is central to the establishment of stable cellular senescence^46,73–76^, we first analyzed their co-expression pattern. As expected, the majority of co-expressing cells were in cluster 2 (Fig. 3E), and were roughly twice as abundant in actively infected and bystander cells compared to mock-infected naive cells (Fig. 3F, Supplementary Fig. 9). This differential abundance was preserved when we analyzed the number of co-expressed genes within each cluster (Fig. 3G, Supplementary Fig. 9). Thus, while the actively HIV-infected cells comprise only 7.48% of cells in the infected (i.e. HIV-exposed) sample, HIV exposure has doubled the overall rate of stable cell cycle arrest in the infected population compared to the HIV-naive cells. Next, we used the list of canonical senescence and SASP genes^69^ which were represented in our dataset (11 genes) to assess the co-expression pattern of the SASP genes with one or both of the hallmark senescence genes expressed in our dataset (Fig. 3H, I, J). We saw up to ten of our 11 genes co-expressed in some cells (Fig. 3H, I, J). Using a minimum co-expression level of 6 senescence/SASP genes, senescent cells comprise 4.63%, 3.13% and 2.17% of actively infected, bystander, and HIV-naive cells, respectively. Comparing the distribution pattern of actively infected and naive cells showed a significant difference (p-value = 0.00011, Fig. 3I). To ensure that the observed patterns were not reflecting the inherent transcriptomic imbalance among cellular populations, we created three control sets by identifying expression-matched gene pools for each senescence gene and randomly drawing one gene per pool to form each control set. Identical co-expression analyses on these control gene lists revealed drastically different patterns, with neither control group showing over 6 co-expressed genes (Supplementary Fig. 10). Further, neither of the control gene sets showed a statistically significant difference between the distribution of actively-infected and naive cells (Supplementary Fig. 10). Thus, after only seven days of HIV infection, the early signs of the development of HIV-induced senescence have developed, and importantly, in addition to actively infected cells, bystander cells also show entry into senescence following HIV infection.

To confirm these results, we used a publicly-available dataset of iPSC-derived microglia grown in triculture with iPSC-derived astrocytes and neurons^47^. This datasets had an infection rate of ∼15%, allowing a similar comparison of actively infected, bystander, and HIV-naive cells as we performed above in our dataset. Importantly, the microglia were infected in monoculture with the CSF-derived, R5-tropic JAGO strain of HIV followed by 15 days of maintenance, before addition to tri-culture for 14 days at which point the cells were harvested for scRNA-seq. Thus, these microglia represent a much later infection timepoint (day 29). Clustering revealed four distinct cellular populations, three of which corresponded to microglia, neurons, and astrocytes, as evidenced by the cluster-specific expression of canonical marker genes for each cell type and close similarities to the cell type-specific expression patterns observed in bulk RNA-seq data (Supplemental Figure 11A-E). The fourth cluster showed a less differentiated expression pattern, consistent with immature astrocytes (Supplemental Figure 11A, F, G). Re-clustering of the microglial population led to identification of subsets corresponding to experimental conditions: the uninfected, untreated microglia; HIV infected ones; and HIV infected microglia treated with Efavirenz (Supplemental Figure 12A). All three conditions showed robust expression of canonical microglial markers (Supplemental Figure 12B).

We next classified the microglial population based on the presence or absence of HIV gene expression, identifying the actively infected microglia as those expressing HIV genes at a level above the maximum number of counts that could be detected in uninfected, untreated population which likely reflected alignment errors. Microglia exposed to HIV but not actively expressing HIV genes, which includes both latently infected and uninfected cells, were labeled bystander cells (Figure 4A). We observed the robust expression of inflammatory cytokines including IL1B in microglia actively infected with HIV (Figure 4B). To test for the presence of the senescence signatures in individual cells from actively infected, bystander and uninfected untreated groups, we calculated an aggregate expression score for the pro-senescence genes in the KEGG cellular senescence pathway (Fig. 4C). We observed significantly higher senescence scores in the actively infected cells than uninfected untreated cells (Figure 4C), and indeed, could show statistically significant co-expression of p21/CDKN1A and p15/CDKN2B with HIV in the same cell (Fig. 4D and E). Importantly, we could show that bystander cells also showed an increase in the expression of senescence genes, albeit less than what was observed in actively infected cells (Fig. 4C). Since the bulk RNA-seq analysis identified the p53 pathway as a key mediator of senescence in HIV-infected microglia, we also calculated aggregate expression scores for the p53 pathway. While the actively infected microglia had the highest expression scores for genes in the p53 pathway, the bystander microglia also showed an elevated expression of p53 pathway genes compared to uninfected cells (Figure 4F). MDM2, a direct target of the p53 pathway and a marker of its activation, showed statistically significant co-expression with HIV in the same cell, providing further proof that HIV directly induces senescence in microglia (Figure 4F, G). Importantly, aggregate scores calculated for the mTOR pathway indicated that both HIV infected and bystander cells show an elevated level of mTOR activity, thus ruling out a p53-mediated entry into senescence^38–42^ (Supplementary Fig. 13).

**Figure 4.**
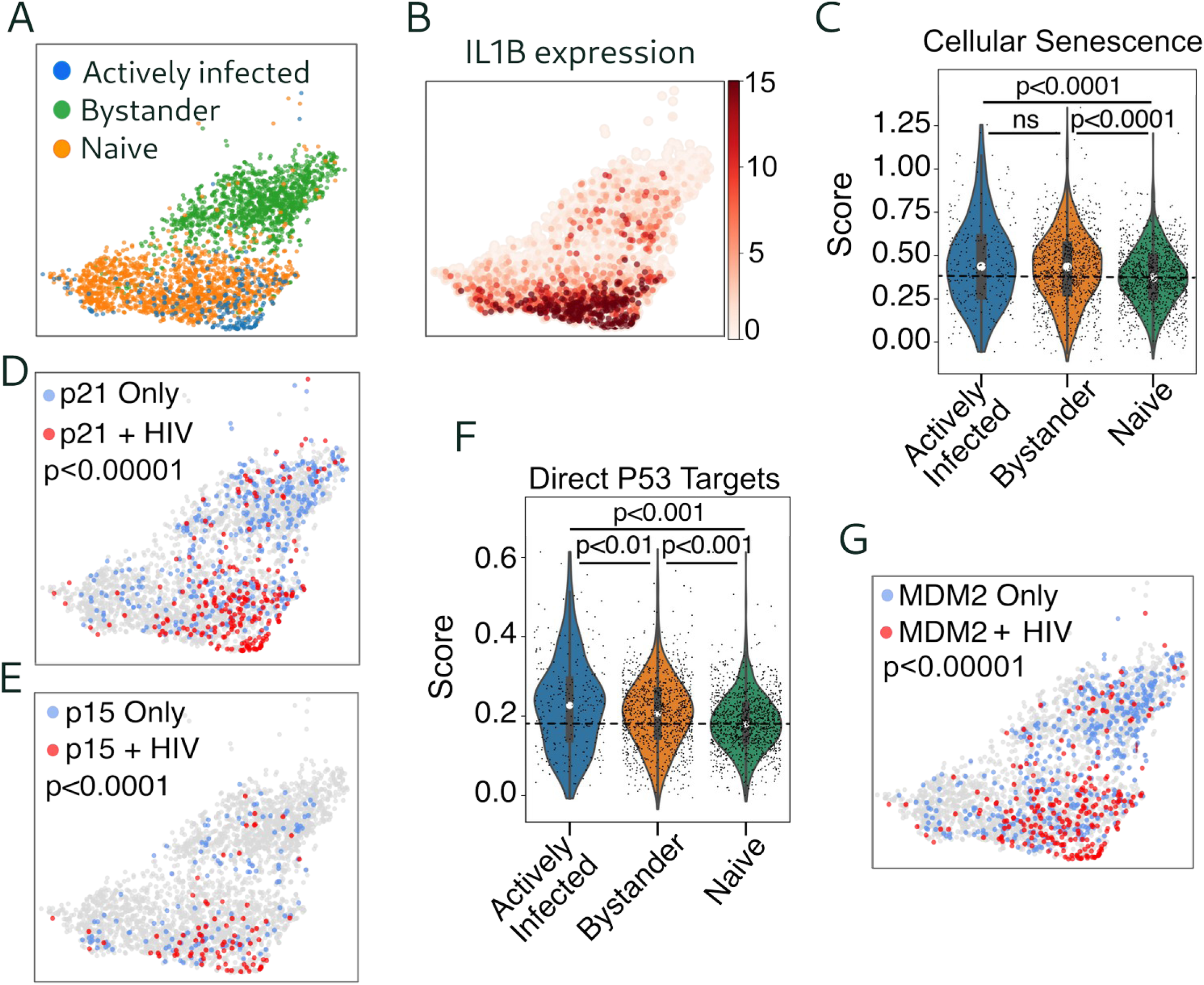
Independent confirmation of induction of senescence in HIV-infected and bystander iPSC-derived micriglial cells. A. UMAP of HIV-infected and uninfected microglia divided into actively infected microglia, bystander microglia, and HIV-naïve microglia. (B) UMAP of IL1B expression in HIV-infected and uninfected microglia. (C) Violin plot of composite expression of genes in the KEGG cellular senescence pathway in actively infected, bystander, and HIV-naïve groups of microglia. (D) Violin plot of composite expression of genes in the direct p53 targets pathway from the C2 data base in actively infected, bystander, and HIV-naïve microglia. (E) UMAP of HIV and p21 co-expression in HIV-infected microglia. (F) UMAP of HIV and p16 co-expression in HIV-infected microglia. (G) UMAP of HIV and MDM2 co-expression in HIV-infected microglia.

### HIV expression drives a cellular trajectory that ends in a state of cellular senescence

We next sought to identify the sequence of cellular states leading to HIV-induced senescence using RNA trajectory analysis on both mock infected HIV-naive and HIV-infected cells as input. Importantly, in these studies the starting and ending states were not specified, allowing the algorithm to deduce the sequence of events in an unbiased manner. Interestingly, the deduced sequence of cellular states started with clusters predominantly containing mock infected, HIV-naive cells (cluster 7) and ended with a cluster almost entirely composed of actively HIV infected cells (cluster 8, Fig. 5A, Supplementary Fig. 14A, B). The intervening clusters represented ordered transition states from uninfected populations to those representing bystander and latently infected cells with low HIV expression levels to actively infected cells (Fig. 5B). We could show that HIV-induced genes identified in our bulk RNA-seq studies were increasing in expression near the end of the trajectory, establishing a direct correlation between viral presence and host transcriptional reprogramming that bridges population-level and individual cellular responses to HIV infection (Fig. 5C). Comparison of the first and last step of the trajectory (clusters 7 and 8, respectively) pointed to the upregulation of inflammatory and senescence markers in cluster 8, and strikingly, to the depletion of expression of neuroprotective genes in this cluster (Fig. 5D, Supplementary Fig. 14C). To further investigate this finding, we leveraged our trajectory to generate a gradient of HIV expression by arranging the clusters along the pseudo-time axis of the trajectory (Fig. 5E, top row), while assessing the expression changes in two neuroprotective genes along the HIV gradient (Fig. 5E). These studies revealed a striking mechanistic relationship between active HIV infection and neuroprotective gene expression profiles. While cluster 1, which is dominated by bystander cells, shows a strong inflammatory and senescence signature which is similar or even higher than cluster 5 (which has a much higher proportion of actively infected cells) (Fig. 5F-L), only cluster 5 shows the depletion of critical neuroprotective factors (Fig. 5E). This suggests that in addition to the known pathogenic impact of cellular senescence and the accompanying inflammatory phenotype, additional factors in actively infected cells play important roles in loss of neuroprotection and the pathogenesis mediated by HIV in the brain. These results further demonstrate the induction of senescence and the accompanying SASP phenotype in bystander cells, as evidenced by the elevated senescence score and expression of CDKN2B/p15, and SASP genes IL-1A and B, IL-6, CCL2 and TNF in cluster 1, which contains ∼70% bystander cells (Fig 5A, F-L). Taken together, our finding provide novel insights into the pathogenic impact of HIV infection in neural tissues. While the strong induction of cellular senescence and the extensive pathogenic reprogramming of genes involved in inflammatory signaling and remodeling of the extracellular matrix is observed in both actively infected and bystander cells, loss of neuroprotective factors were only observed in actively infected cells.

**Figure 5.**
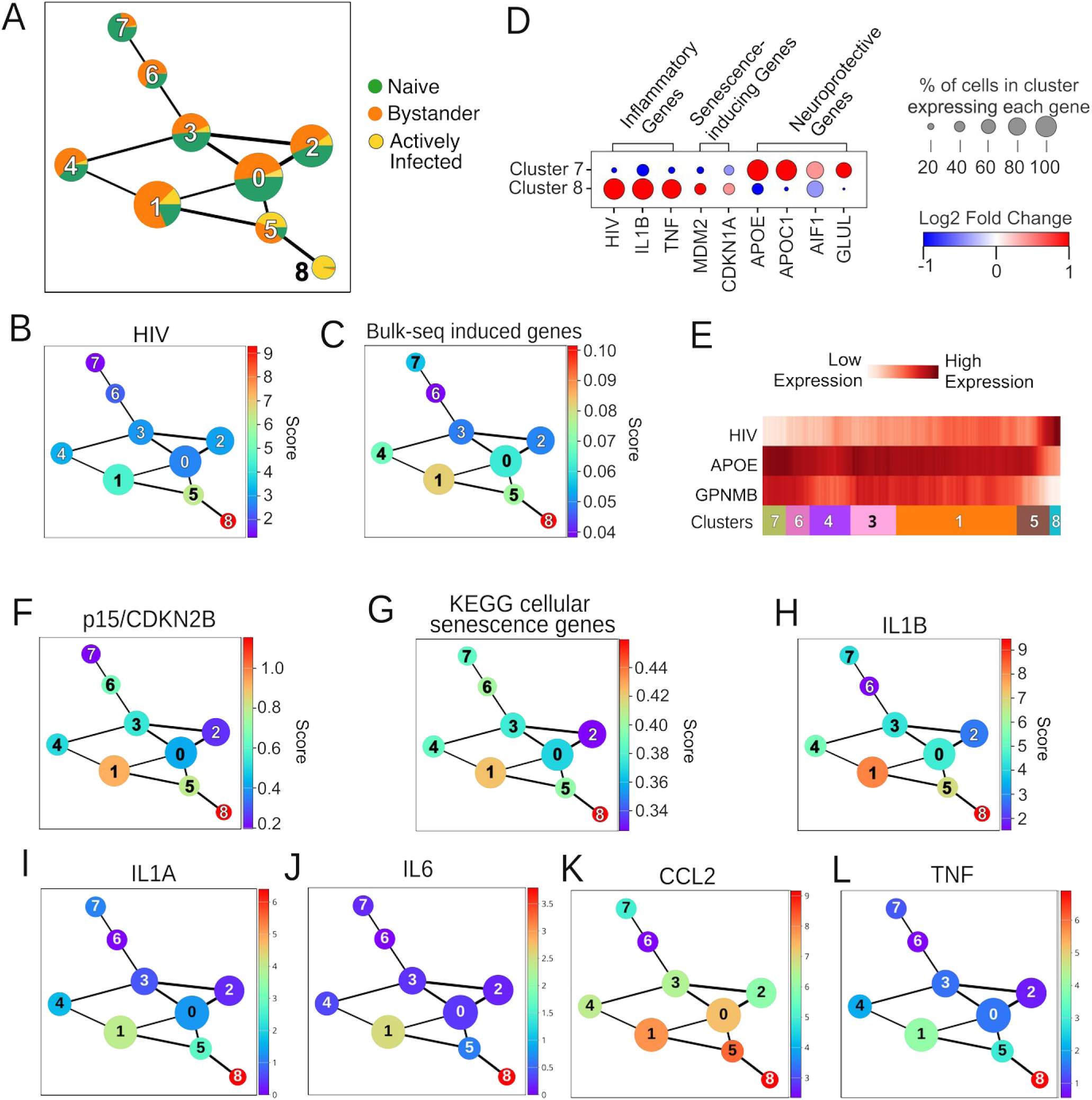
Trajectory analysis reveals the association of progression of senescence phenotype with loss of expression of neuroprotective genes. (A) Percentage of actively infected, bystander, and HIV-naïve microglia in each node of RNA trajectory analysis. (B,C) HIV and TNF expr4ession per trajectory node. (D) Composite expression of genes identified in bulk RNA-seq analysis as increasing in microglia in response to HIV. (E) p15 expression per node. (F) Composite expression of genes in the KEGG cellular senescence pathway per node. (G) Trajectory path plot showing per-cell expression along trajectory of HIV, TNF, APOE, and GPNMB.

### Microglia from in vivo SIV-infected rhesus macaques show the senescence phenotype

While our data indicate that the induction of the dual axes of NFkB and p53 induces senescence in iPSC-derived ex-vivo HIV infected microglia, it is important to validate these findings in a clinically-relevant context. The brain’s microenvironment provides a sophisticated network of cell-cell interactions and regulatory mechanisms which are only partially recapitulated in tri-culture systems but can strongly impact microglial response to HIV infection^77,78^. In addition, the chronic nature of HIV-associated neuroinflammation in patients is not effectively modeled in ex vivo systems, necessitating validation studies using in vivo models.

To this end, we took advantage of the availability of single cell RNA-seq datasets from in vivo SIV-infected rhesus macaque monkeys^79,80^. The study utilized 12 male rhesus macaques (3-7 years old) which were divided into four experimental groups of 3 animals each: uninfected controls, SIV-infected without c-ART treatment, SIV-infected with c-ART, and SIV-infected developing encephalitis (SIVE), the latter representing the relevant model for studying HIV-associated neurocognitive disorders (HAND) in humans^81–83^. The SIVE and untreated SIV groups received an intravenous injection of in vivo (rhesus monkey) passaged SIVmac251 stock, while the SIV-cART group received an in vitro (CEMx174 cells) passaged stock. The cART-treated animals received daily subcutaneous injections of emtricitabine, tenofovir, and dolutegravir beginning 5 weeks post-infection and continuing until necropsy. Animals were sacrificed at different timepoints based on the treatment received, from 65 and 92 days post-infection (for the SIVE animals) to after at least 30 weeks of treatment for the c-ART-treated animals. Half of the brain of the animals were harvested and enriched for myeloid cells using CD11b enrichment followed by scRNA-seq7^9,80^. We used clustering to distinguish the microglial cells from the CD11b+ lymphoid and macrophage-like cells using lineage markers to identify each cluster (Fig. 6A, Supplementary Fig. 15, S16). Analysis of the expression pattern of senescence markers pointed to a subset of cells in cluster 6 from animals with SIVE, which consistently showed the expression of senescence markers and genes involved in SASP (Supplementary Fig. 17). Importantly, this cluster also had a high density of SIV+ cells (Supplementary Fig. 17).

**Figure 6.**
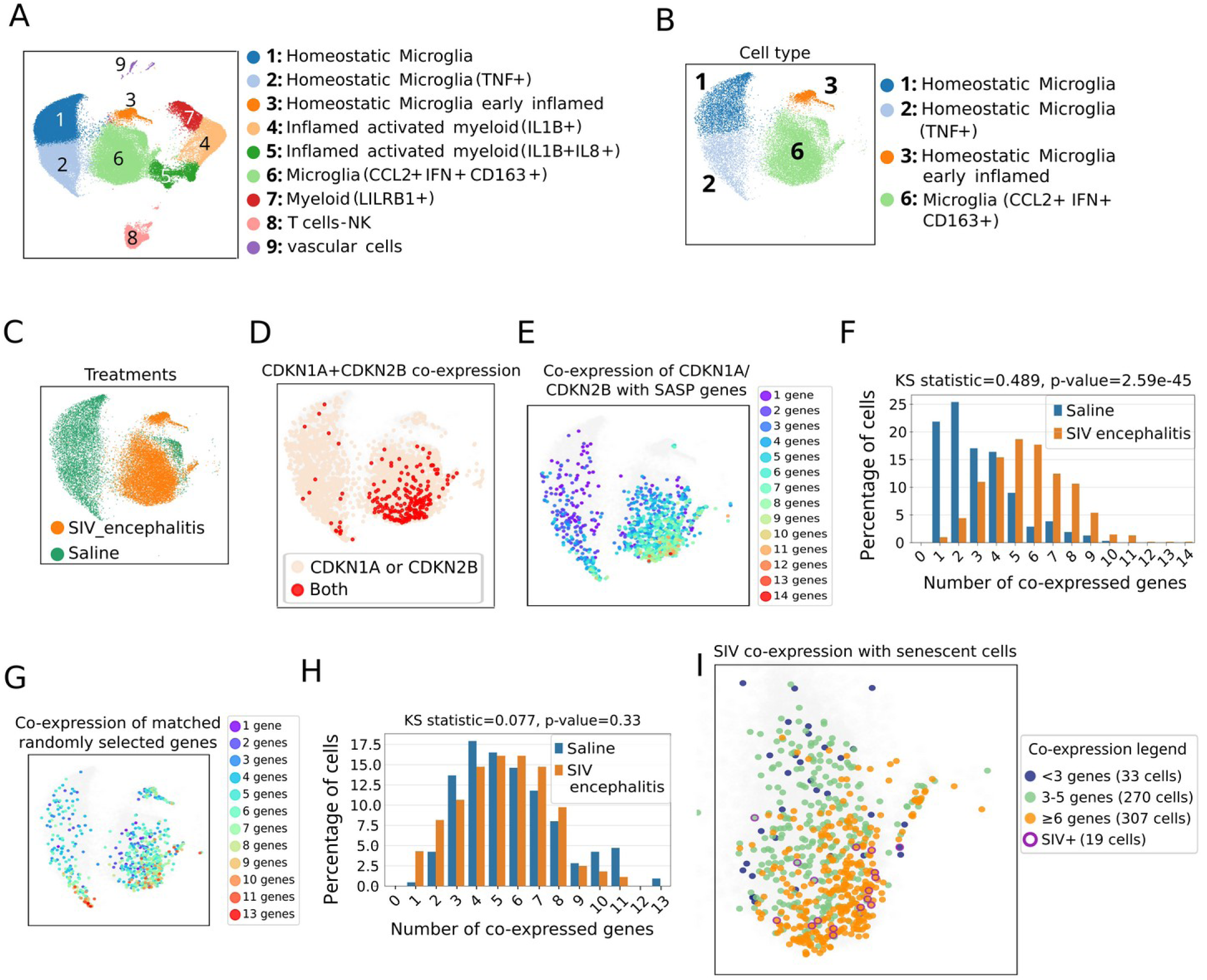
Induction of cellular senescence in SIVE animals. A. UMAP of CD11B+ cells purified from the brains of saline-treated or SIV-infected animals, with and without c-ART. B. UMAP of microglial states represented in saline-treated and SIV-encephalitis (SIVE) animals in the absence of c-ART. C. UMAP indicating the distribution of cells from each treatement group. D. Co-expression of p21/CDKN1A and p15/CDKN2B is strongly enriched in SIVE animals. Red dots mark cells co-expressing both senescence markers. E. Co-expression pattern of markers of senescence and SASP, representing senescent cells. Colors corresponding to the number of coexpressed senescence and SASP markers in each cell are indicated. F. Barplot of the percentage of cells in saline-treated and SIVE animals showing each level of co-expression of senescence and SASP markers. The provided p-value corresponds to the comparison of the overall co-expression pattern of saline and SIVE samples. G and H. Similar analyses to E and F, using a set of control genes selected to match the expression pattern of the senescence and SASP markers (see Methods). I. Overlap of actively infected and senescent cells. We used a threshold of 6 coexpressed genes for defining the senescent cell population.

We focused on the microglial population from the saline-treated and SIVE samples, which includes the cluster with high levels of senescence-related genes in addition to constituting the most relevant model for the study of HAND (Fig. 6B, C, Supplementary Fig. 15B, C, F). We first analyzed microglia for expressing senescence markers along with a panel of SASP-related genes, followed by comparison of the expression levels between the SIVE and saline-treated samples (Supplementary Fig. 17, 18, see Methods). We compared the percentage of cells in each treatment group which were positive for each gene, along with the mean expression level of each gene among cells which expressed the gene. To identify statistical significance, we employed generalized linear mixed models (GLMMs) to control for transcriptome size variation between experimental groups, ensuring robust detection of differential expression patterns. We used a hurdle approach which allowed us to integrate the two gene expression metrics (see Methods). The majority of our marker genes showed a strong and statistically significant enrichment in cells from SIV animals compared to saline-treated ones.

To identify senescent cells, we used an approach similar to the one used for the iPSC-derived microglia, namely, testing for the combined presence of two primary senescence signature genes (i.e. p21/CDKN1A, p15/CDKN2B or p16/CDKN2A), or the presence of at least one of them along with several supplementary markers (including p27/CDKN1B, p62/SQSTM1 and SASP-related genes)^69–72^. We first analyzed the pattern of co-expression of the primary senescence marker genes. The co-expression of p21/CDKN1A and p15/CDKN2B, both hallmark senescence markers, is indicative of a stable, long-term block to cell cycle and a strong indicator of entry into senescence^46,73–76^.

Their co-expression was observed in over 4 times as many cells in SIVE than saline samples (251 (1.11%) vs. 56 (0.25%) cells, p-value 1.79e-08 after adjusting for transcriptome size, see Methods), indicative of accelerated microglial senescence in SIVE animals (Fig. 6D). To take the high rate of dropouts in single cell RNA-seq into account, we also identified cells which expressed a combination of either p21/CDKN1A or p15/CDKN2B, along with other genes from a widely accepted list of 19 senescence- and SASP-associated genes^69^ (see Methods). We observed a strong pattern of co-expression of senescence markers and SASP-related genes in SIVE samples, while saline-treated samples showed a markedly weaker co-expression, consistence with a much higher proportion of cells showing robust senescence phenotype in SIVE animals (Figs. 6E, F, Supplementary Fig. 19). Similar results were observed when individual study subjects in the two experimental groups were compared (Supplementary Fig. 20) Comparison of the co-expression patterns of the two groups using Kolmogorov– Smirnov (KS) test showed a large effect size (KS statistic of 0.489, p-value=2.59e-45, Fig. 6F). To ensure that the observed difference between SIVE and saline-treated samples did not stem from technical artifacts such as differences in dropout rate or transcriptome size, we performed parallel co-expression analyses using four sets of control genes with closely similar gene expression patterns to the senescence and SASP genes used in our study, as used in the analysis of iPSC-derived microglia (see Methods). The co-expression pattern of these randomly selected genes were largely similar between SIVE and saline-treated samples, with a small effect size (KS statistics ranging from 0.075-0.095) and non-significant p-values (Fig. 6G, H, Supplementary Fig. 21). Similarity of the results from multiple random gene sets established a baseline level of co-expression in our samples, against which the senescence-associated co-expression pattern could be meaningfully evaluated. Overall, 307 (2.2%) of microglial cells in the SIVE samples showed the co-expression of at least 6 senescence and SASP-related genes (including either CDKN1A or CDKN2B), compared to 32 (0.3%) in saline samples, a difference of over six fold which provides proof for the strong SIV-mediated induction of senescence in microglia in vivo. The small number of senescent cells in saline samples likely reflect aging-associated senescence. Due to the high dropout rate in scRNA-seq, the co-expression values are often strongly underestimated, and thus, the true number of senescent cells in SIVE samples is likely to be much higher than estimated using scRNA-seq. Nonetheless, comparison of co-expression values between SIVE and saline-treated samples, which are of similar transcriptome size range and composition (Supplementary Fig. 16, 21) is a strong indicator of the relative co-expression patterns.

While our studies in ex vivo iPSC-microglial tri-cultures show the strong induction of senescence pathways in actively infected cells, the presence of the full immunological milieu and brain microenvironment is likely to impact the senescence dynamics. For example, immune response can clear the actively infected cells before they develop a senescence phenotype or strongly restrict the proviral replication, thus reducing the pool of actively infected cells. So, in order to fully understand the dynamics of SIV-induced senescence in vivo, we identified the actively infected cells (defined as cells with detectable expression of SIV, Fig. S17). Analysis of the overlap of SIV+ cells and senescent cells showed an increased association with senescent cells (those co-expressing 6 or more senescence-related genes) compared to cells not reaching this threshold (Chi-square statistic 4.523, p-value 0.033, Fig. 6I). This finding confirms the enhanced senescence phenotype in actively infected cells that was observed in ex vivo iPSC-derived models. However, the lower magnitude of the effect, which was also confirmed by calculation of aggregate scores for expression of pro- and anti-senescence genes in total bystander and actively infected population (Supplementary Fig. 22), likely point to important differences between the in vivo and ex vivo models. We performed a differential expression test between SIV+ and bystander cells, which did not show any significant gene expression changes except for the SIV gene itself. Comparison of SIV+ or bystander cells to saline-treated cells yielded over 1600 differentially expressed genes (>2 fold change, FDR<0.05), which were shared between the two comparisons. These results suggests that the overall pattern of gene expression of actively infected and bystander cells are largely similar in vivo.

### Treatment with c-ART reverses most, but not all, of the SIV-induced senescence signatures

While SIVE samples are the most relevant model for the study of HAND, an important remaining question is whether SIV infection in the absence of encephalitis can also induce senescence. Further, it is important to determine the impact of cART treatment on senescence. To this end, first asked if SIV-infected samples (non-encephalitis, not treated with cART) show a senescence signature when compared with mock-infected samples. No SIV-infected cells were detectable in these samples, likely reflecting the immune clearance of infected cells and entry of the remaining infected cells into latency. Analysis of the co-expression of CDKN1A and CDKN2B indicated that SIV infection was associated with a mild increase in the fraction of double positive cells (1.0% versus 0.6% for saline-treated samples, Fig. 7A, B). Following correction for transcriptome size, this difference corresponds to a significantly increased odds of co-expression of the two genes in SIV samples (Odds Ratio=5.76 [95% Confidence Interval : 3.90–8.51], p-value<<2e-16). We also determined the extent of co-expression of one or both of CDKN1A and CDKN2A with SASP genes. These studies revealed a clear increase in representation of cells with higher co-expression levels (>=6 co-expressed genes) in SIV samples relative to saline (2.27% vs 0.36%, respectively, p-value 2.9e-4, Fig. 7C, D). Four sets of gene sets showed distinct co-expression distribution patterns, none of which was statistically significant (Supplementary Fig. 23). Thus, SIV infection in the absence of encephalitis does induce a moderate increase in the number of senescent cells. As the actively infected cells are not detectable in cells from either of these conditions, the increase in senescence-prone cells correspond to the impact on bystander cells.

**Figure 7.**
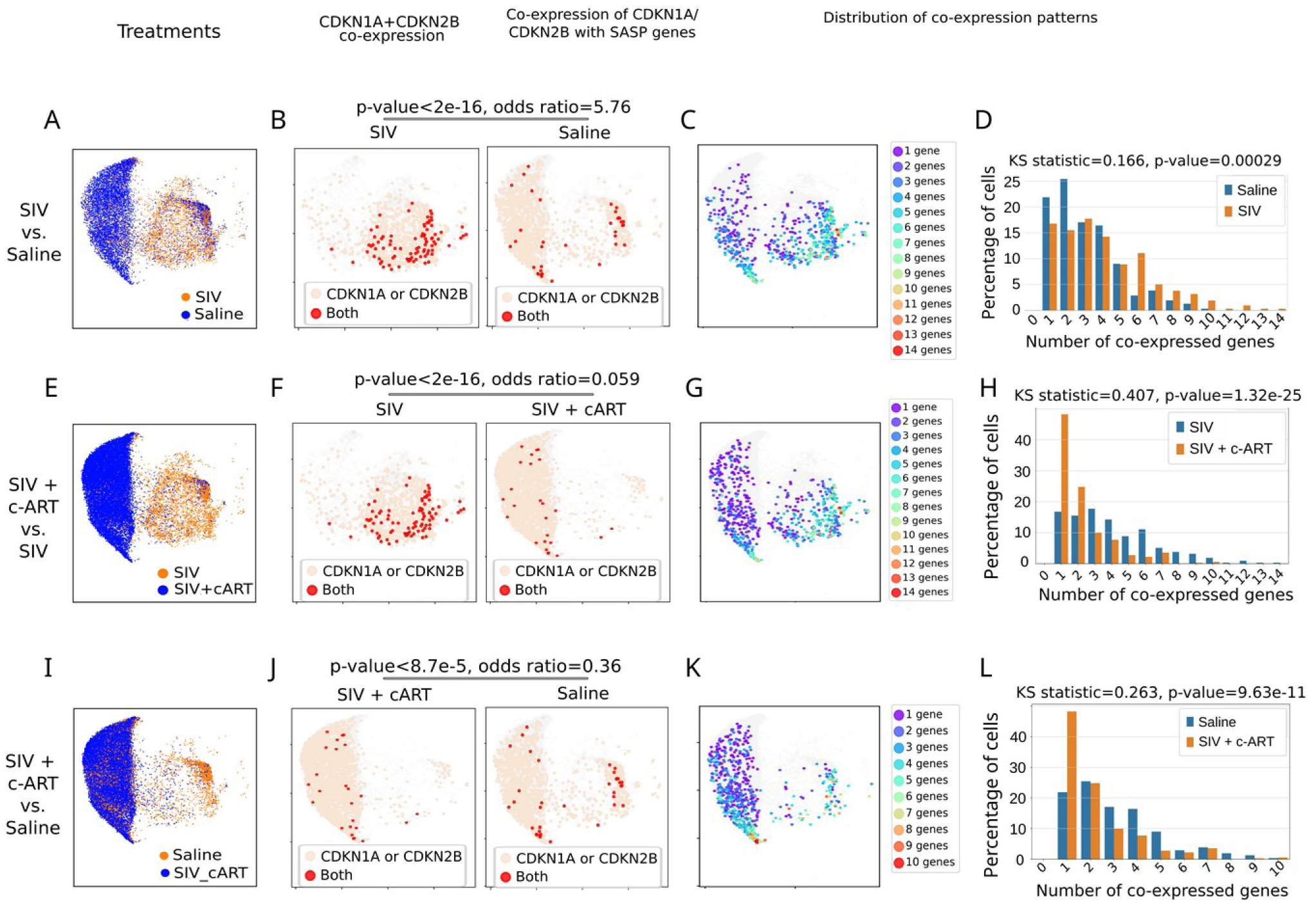
c-ART treatment in SIV-infected cells largely reverses the SIV-induced expression of senescence and inflammatory genes. Pairwise comparison of SIV-exposed microglia with saline-treated (A-D), SIV+cART versus SIV (E-H) and SIV+cART vs saline-treated samples (I-L) are shown. For each comparison, the co-expression level of CDKN1A/p21 and CDKN2B/p15, and that of senescence markers and SASP genes are shown. Quantitation of the co-expression results along with the statistical test results indicates the presence of small, but significant changes induced by SIV infection and cART treatment (D, H and L).

To determine if treatment with cART reverses the SIV-induced senescence patterns, we compared the senescent cellular fraction in the two conditions. In SIV and SIV+cART samples, 1.01% and 0.12% of cells co-expressed CDKN1A and CDKN2B, respectively, corresponding to an odds ratio of 0.059 and a p-value of < 2×e-16 (Fig. 7E, F). Comparison of the co-expression of senescence markers with SASP genes showed a statistically significance difference between the two groups, with 2.27% and 0.121% of SIV and SIV+cART cells co-expressing at least 6 genes (p-value 1.32e-25, Fig. 7G, H, Supplementary Fig. 24). This significant drop in senescent cells indicates that cART can reverse the induction of senescence genes in animals infected with SIV. Finally, the difference in co-expression of senescence markers between SIV+cART and saline-treated samples remained statistically significant (Fig. 7I-L, supplementary Fig. 25), suggesting that cART-treated cells are not fully restored to the homeostatic state. Analysis of the aggregate expression pattern of inflammatory response genes and p53 pathway activation pointed to an ‘under-inflamed’ or hypo-responsive state with reduced expression of p53 pathway and inflammatory genes relative to saline-treated microglia (supplementary Fig. 26A, B). As a complementary approach, we calculated the aggregate score for pro- and anti-senescence genes from the CellAge database^84^. Surprisingly, we observed changes in SIV + cART samples which mimicked those in SIVE and SIV samples, but were absent in saline-treated, homeostatic microglia (Supplementary Fig. 26C, D). These results in aggregate suggest that while as previously reported c-ART largely reverses HIV and SIV-induced pro-senescence effects^24,50,57,79,85^, some residual senescence-related changes and a hypo-responsive state may impact the microglial function during cART. This, in turn, may contribute to the asymptomatic and mild cognitive impairments seen in PLWH who are also on a cART regimen.

### Microglial senescence is associated with the induction of neurotoxicity pathways

In addition to the induction of senescence and its associated inflammatory factors and remodeling of the extracellular matrix, HIV/SIV infection may induce additional functional changes in microglia that contribute to HAND. As immune cells, microglia largely mediate their function through cell-to-cell communication via secreted and membrane-bound factors including ligand-receptor interactions. To systematically analyze the functional change in microglia in the context of HIV/SIV infection, we analyzed the ligand-receptor-mediated interactions involving microglia in actively infected, bystander, cells infected with HIV and treated with cART, and SIV-naive, mock infected controls. We used clustering to further group microglia based on their level of inflammatory signaling into resting, homeostatic, activated and highly inflamed to define the interactions involving each group. For these studies, we also included the highly inflamed cells which likely contain non-microglial myeloids cells (clusters 4, 5 and 7 which we had excluded from our other analyses, Fig. 6A) as their signaling activity can impact the pathogenesis of HAND in important ways.

Using the CellphoneDB tool ^86^, we identified receptor-ligand interactions which were differentially enriched between the three groups exposed to HIV and the mock-infected, SIV naive cells (Fig. 8A). We identified over 40 differentially enriched interactions. Among them were several ligand-receptor interactions known to enforce a durable G₁-phase arrest and fuel SASP in SIV-infected microglia (Fig. 8B-D). The Prostaglandin E₂:PTGES2:PTGER4 pathway, which sustains autocrine inflammatory feed-forward loops in senescent cells^87^, was selectively upregulated in actively infected and bystander microglia and returned toward baseline in cART-treated cultures (Fig. 8B-H). IL-15:IL-15R engagement and TWEAK (TNFSF12):TNFRSF25 signaling, both implicated in NFκB-dependent SASP gene expression, were similarly upregulated in actively infected and bystander cells, and interestingly, also showing strong activity in SIV+cART cells (Fig. 8B-D, Supplementary Fig. 27). Interestingly, TGF-β1:TGFβR1/2/3 signaling, which drives p15/p21 induction and Rb-mediated cell-cycle exit^88^, remained active in the SIV + cART group and is even elevated in homeostatic microglia relative to actively infected and bystander groups, indicating that cART cannot fully restore homeostatic state in cells infected with/exposed to SIV (Fig. 8B-D). Together, these data confirm that SIV infection initiates a robust senescence program in microglia that persists, though at reduced amplitude, following anti-retroviral treatment.

**Figure 8.**
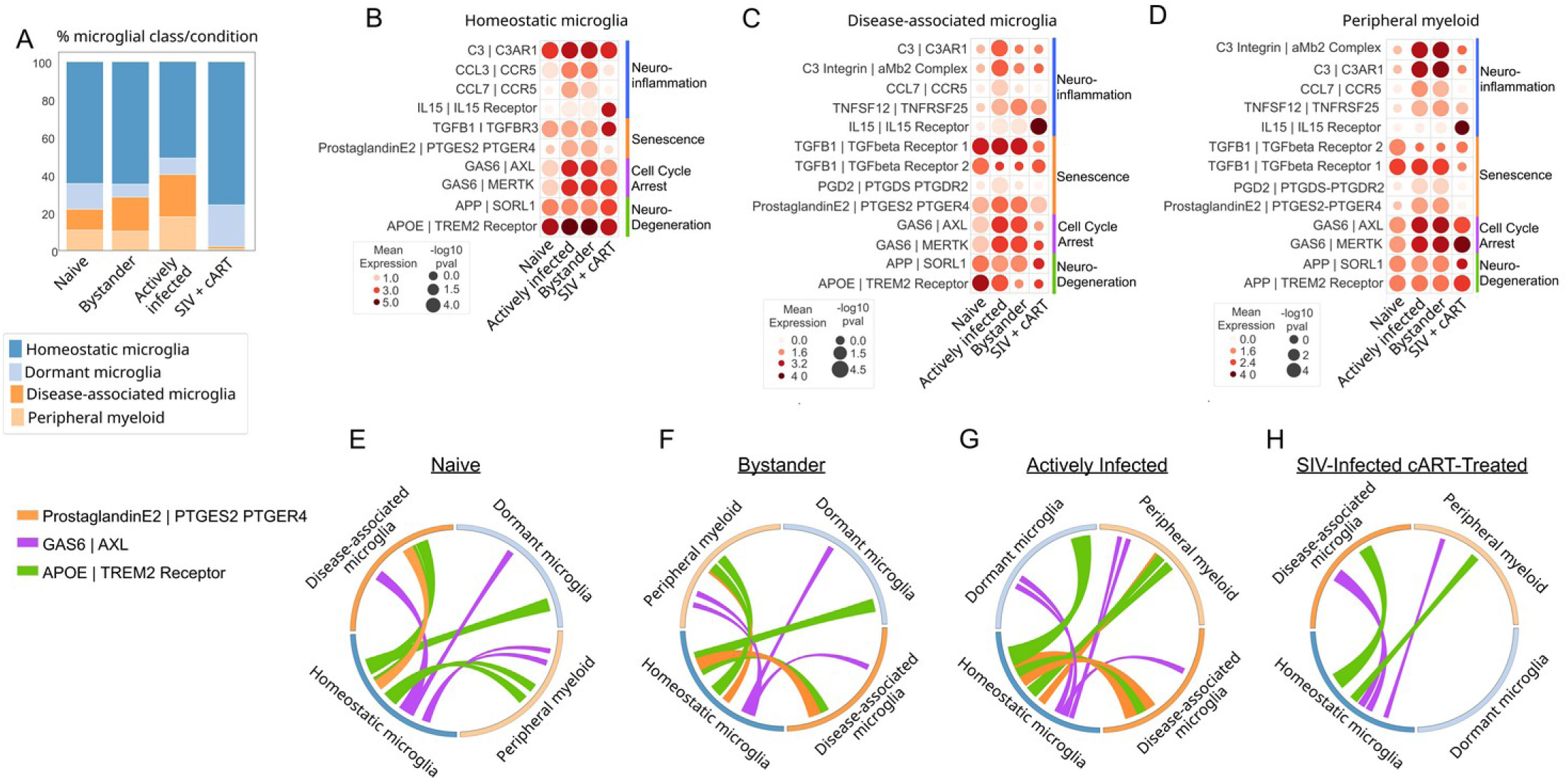
Microglia from Mock-infected (naive), SIV-infected, and SIV-infected cART-treated (SIV + cART) Macaques show changes in senescence and neuro-inflammatory signaling. (A) Barplot of relative abundance of different microglial classes per condition. (B,C,D) Dotplots showing the mean level of outgoing signals from homeostatic microglia, disease-associated microglia and peripheral myeloid cells. (E,F,G,H) Chord plots of ligand-receptor pairs per condition.

A second group of enriched interactions involves neurotoxicity and neurodegeneration pathways. C3:C3AR1, iC3b:αMβ2 (Mac-1) integrin, and CCL3/CCL7→CCR5 chemokine signaling were among the top enriched pathways in actively infected and bystander microglia, declining sharply under cART (Fig. 8B-D). Thus, active SIV exposure drives a classic neurotoxic signature in microglia characterized by complement and chemokine axes that promote synaptic pruning and immune cell recruitment^89^.

These signaling patterns reveal a complex microglial response to SIV infection characterized by two key mechanisms: activation of senescence pathways and establishment of neuroinflammatory pathways that partially persist despite cART treatment. The coordinated activation of both senescence and neuroinflammatory pathways supports our earlier findings (Fig. 5) that these processes are mechanistically linked in SIV-infected microglia and contributes to the pathophysiology of HAND. Interestingly, the strongest changes in signaling patterns compared to uninfected, saline-treated naive microglia were observed in cART-treated samples, which showed sub-homeostatic signaling in multiple functionally critical cell-cell interactions (Fig. 8B-H). In agreement with our earlier analyses, these results point to a hypo-reactive, under-inflamed phenotype in cART-treated cells.

## Discussion

Our analysis of three ex-vivo and one in vivo model for HIV/SIV infection in the brain, we have been able to show that HIV/SIV induce a strong, multi-pronged senescence program in microglia, leading to the induction of multiple neuroinflammatory pathways and loss of expression of neuroprotective genes. Importantly, we were able to prove that in addition to cells actively expressing HIV/SIV genes, a sizable fraction of bystander cells, which are either not infected or harbor a latent virus, also show a strong senescence phenotype. This, in turn, leads to tissue damage and through secretion of inflammatory cytokines, likely triggers a vicious cycle of senescence induction. Importantly, we were able to directly compare HIV-induced changes in microglia with those resulting from organismal aging, showing that both pathways converge on the same cellular pathways and thus, show synergy in induction of cellular senescence and SASP, thus providing a mechanism for the HIV-induced exacerbation of the impact of aging.

Our use of multiple datasets including three independently performed ones and an in vivo study proves the reproducibility and generality of our results. Further, as the used datasets corresponded to different timepoints after HIV infection (7, 15, 29 and >60 days post infection), the differences in the level of induction of senescence, which was lowest, but nonetheless clearly present, in the 7 day post-infection sample and highest in the in vivo, >60 days post-infection one, could point to the natural history of HIV-induced senescence. It is known that acute HIV infection leads to activation of the infected cells, while at the same time triggering a strong inflammatory response and oxidative, ER and mitochondrial stress which activate the p53 pathway. The combination of cellular activation, HIV integration and dual induction of NFkB and p53 pathways can initiate the activation of pathways involved in engagement of the RB axis and thus, initiate the cellular senescence process. Interestingly, in our earliest timepoint (7 days post-infection sample), we observed that the cluster showing the induction of p16/CDKN2A which engages the RB axis also expressed proliferative genes (Supplementary Fig. 28), potentially capturing the transition of the induction of cellular activation by HIV toward a cell cycle arrested senescence state. Our findings support a model where HIV infection initially triggers a mixed activation/proliferation response that gradually resolves toward senescence as cellular stresses accumulate and p16-mediated cell cycle arrest becomes dominant. Importantly, the impact of co-expression of p21/p16 extends beyond mere cell cycle regulation, as these proteins participate in broader cellular processes including chromatin remodeling, DNA damage response, and the modulation of SASP, collectively orchestrating the complex physiological state of cellular senescence that influences tissue homeostasis, aging, and disease progress^46,73–76^.

An interesting aspect of the emergence of cellular senescence is its impact on HIV life cycle. The relationship between HIV/SIV infection and cellular senescence reveals a nuanced interplay of host defense and viral adaptation. While senescence initially functions as a protective cellular response that limits viral replication, HIV has developed sophisticated mechanisms to repurpose aspects of this process for its benefit, notably exploiting the increased transcriptional environment during G2 cell cycle arrest to enhance proviral integration. Viral proteins contribute significantly to this phenomenon, with Tat promoting senescence through NFκB pathway activation and Nef through autophagy inhibition. These proteins are released from infected cells and induce senescence in neighboring uninfected cells, creating a microenvironment that simultaneously restricts new infection while optimizing conditions for already established viral reservoirs. This dual effect likely represents an evolutionary balance between viral persistence and limitation of pathogenicity^90^.

Interestingly, multiple lines of evidence in our studies including trajectory and cell-cell interactions pointed to a coordinated induction of senescence and neuroinflammatory pathways, together with loss of neuroprotective ones. The elevated level of neurotoxic signaling together with microglial malfunction caused by senescence and the inflammatory environment, can largely account for the pathogenesis of HAND. This, in turn, suggest that interventions aimed at mitigating microglial senescence and restoring homeostasis could have dual benefits for addressing HAND and age-related neurocognitive decline.

In conclusion, our study highlights the striking parallels between HIV-induced and aging-associated changes in microglia, providing new insights into the cellular mechanisms underlying HAND. These findings lay the groundwork for future research aimed at developing targeted therapies that address both HIV-related and age-related neurocognitive decline.

## Resource availability

The transcriptomic dataset generated by our team for this study will be deposited in SRA and GEO. Other used datasets are available at SRA and GEO repositories under the following accession numbers: GSE143685, GSE99074, GSE111972. No original code was generated during this study.

## Acknowledgments

We gratefully acknowledge the National Institutes of Health awards P30 DA054557 to A.D.L.; R01 MH134316, R01 DA060489, R01 DA049481 and P30 AI036219 (CFAR) to J.K., and NIH HIV Cure Training Grant 5T32AI127201-05 to S.J.M. We acknowledge members of Valadkhan and Karn laboratories for providing insights and sharing resources.

## Author contributions

Conceptualization, S.J.M., S.V., A.D.L. and J.K.; Investigation, S.J.M., S.V., S.S., K.L., A.D.L. and J.K; Dataset preparation, K.L. and S.S, funding acquisition, S.V., J.K. and A.D.L.; supervision, S.V.; formal analysis, S.J.M., F.N.S.; visualization, S.J.M., F.N.S. S.V.; writing – original draft, S.J.M.; writing – review & editing, S.J.M., S.S., K.L., S.V., A.D.L. and J.K.

## Supplemental information

Document S1. Figures S1–S28

## Methods

List of computational resources used in this study

| REAGENT or RE-SOURCE | SOURCE | IDENTIFIER |
| --- | --- | --- |
| Deposited data |  |  |
| iPSC-derived human microglia infected with HIV or mock infected | Bulk and Single cell RNA-seq, Ryan et al. | GSE143685, [10.1016/j.stemcr.2020.02.010](https://doi.org/10.1016/j.stemcr.2020.02.010) |
| Aged microglia from post-mortem brain samples from otherwise healthy donors, parietal cortex | Bulk RNA-seq, Galatro et al. | GSE99074, [10.1038/nn.4597](https://doi.org/10.1038/nn.4597) |
| Aged microglia from post-mortem brain samples from otherwise healthy donors, corpus callosum and occipital cortex | Bulk RNA-seq, van der Poel et al. | GSE111972, [10.1038/s41467-019-08976-7](https://doi.org/10.1038/s41467-019-08976-7) |
| Rhesus macaques either mock infected or infected with SIV, with and without cART treatment | Single cell RNA-seq, Fox et al. | GSE195574, [10.3389/fimmu.2022.1012884](https://doi.org/10.3389/fimmu.2022.1012884) |
| Human Aging Genomic Resource: CellAge | Avelar et al. | https://genomics.senescence.info/ |
| MSigDB | Subramanian et al. | https://www.gsea-msigdb.org/gsea/msigdb |
| Software and algorithms |  |  |
| Kallisto | Bray et al. | https://pachterlab.github.io/kallisto/about |
| Sleuth | Pimentel et al. | https://pachterlab.github.io/sleuth/about |
| GSEA | Subramanian et al | https://www.gsea-msigdb.org/gsea/index.jsp |
| scverse: SCANPY, Ann-data, PAGA (trajectory analysis) | Virshup et al. | https://scverse.org/ |
| Scikit machine learning | Pedregosa et al. | https://scikit-learn.org/stable/index.html |
| CellphoneDB | Vento-Tormo et al. | https://www.cellphonedb.org/ |

## Method Details

### Preparation of iPSC-derived microglial cells

#### Generation and maintenance of tdTomato-tagged iPSC lines

The iPSC line used in this study was generated from a neurotypical male individual (“Clay” from Marchetto et.al ^1^). Reprogramming and validation of this cell line was previously described in Marchetto et.al 2016 and is also described in Bury et. al 2023^2^. All iPSC lines were validated for pluripotency markers OCT3/4, and confirmed normal karyotype. iPSC lines were maintained in complete TeSR-E8 medium (Stem Cell Technologies; Catalog # 05990) until confluency and detached as small aggregates using ReLeSR (Stem Cell Technologies). The aggregated were centrifuged at 1000rpm for 1 minute. The pellet was resuspended in stem cell medium plus rock inhibitor Y-27632 (RI-10 mM) and plated on 6-well tissue culture plates coated with geltrex (Thermo fisher). The cells were incubated at 37°C in a 5% CO2 incubator. Geltrex-coating was performed by mixing geltrex aliquots thawed on ice with ice cold DMEM medium at 1:100 dilution followed by incubating 1ml/well for 1h at 37°C in 5% CO2.

#### Generation of hematopoietic stem cells (iHSC) and microglia (iMGs) from iPSCs

To generate iHSC, iPSCs were differentiated to hematopoietic stem cells using the STEMdiff hematopoietic kit (Stem Cell Technologies). Briefly, iPSCs were dissociated using ReleSR into small aggregates of sizes between 100um-150um and plated at ∼80-100 aggregates/well in a 6 well plate in E8 media with RI. The total colonies plated were carefully selected for each iPSC line post testing different dilutions of colonies per well and yields of CD34+ cells. After confirming for optimal size and number of colonies next day (10-20 colonies/cm^2^), E8 media was replaced with fresh media A at day 0 and 2 of differentiation. To induce the second phase of cell differentiations, media A were replaced with media B at day 3. The cells were given fresh media B every alternative day starting from day 5 to day 12. Beginning day 7, we began seeing round and floating iHSC. On day 12 floating iHSC were harvested by gentle pipetting and characterized for hematopoietic stem cell markers CD43, CD34 and CD45 by flow cytometry.

To induce iMG differentiation from iHSC, we followed the protocol by Blurton-Jones group ^3^ and transferred the day 12 iHSC to iMG media (as described in Abud et.al ^4^) containing 100 ng/mL IL-34, 50ng/mL TGFβ1 and 25ng/mL M-CSF (Invitrogen). iMGs were cultured for upto Day 35 in iMG media, with addition of maturation factors CD200 and CX3CL1 at day 25 until day 35 for obtaining fully differentiated iMGs. We validated iMG surface protein markers by flow cytometry prior to performing scRNA-seq analysis as described later.

#### HIV-1 preparation and infection of iMGs

To generate infectious virions of the replication-competent, macrophage-tropic HIV-1 NL-AD8 strain, we transfected 293T cells in 6-well plates with 4 µg/well of the pNL-AD8 plasmid (Schubert et al., 1999). To enhance viral titers, we used culture supernatants collected 2 days post-transfection of 293T to infect a human monocytic cell line engineered to express human CCR5 and a GFP reporter under the control of the HIV-1 5’ LTR promoter. The infected monocytic cells were then cultured in RPMI 1640 medium supplemented with 10% FBS for 10 days. Cells were collected by centrifugation (1,500 rpm, 3 minutes), resuspended in microglia media, and cultured for an additional 2 days. The final culture supernatant was harvested and used to infect the iMGs

To induce HIV infection of iMGs, we introduced replication competent NL-AD8 directly into cell culture media at day 27 of iMG culture in a 24-well plates. We continued to monitor the cultures for up to 7 days post-infection (dpi) and added additional media every alternate day.

#### Data processing: bulk RNA-seq

Raw bulk RNA-sequencing data from microglia samples from three publicly available datasets (GSE143685, GSE99074, GSE111972) were downloaded, and quality was assessed using FastQC (Babraham Bioinformatics). The reads were pseudo-aligned to the transcriptome and quantified using kallisto 0.46.0, followed by normalization using sleuth^91,92^.

Differential gene expression analysis was performed, and genes with over 2-fold changes in expression and an FDR of 0.05 or lower were used for pathway analysis. Pathway analysis was conducted using GSEA version 4.2.1^93^ with version 7.5.1 of the Hallmark pathway geneset from MSigDB and the KEGG database ^94–96^. Pathways with an FDR q-value ≤ 0.05 were considered significant.

Correlation analysis was performed using Spearman and Pearson coefficient tests. The coefficients for each gene were averaged, and genes with an averaged correlation coefficient greater than 0.4 or less than −0.4 were selected for pathway analysis, which was conducted as described above.

#### Single cell RNA-seq analyses

Raw single cell RNA-sequencing data from microglia samples from two publicly available datasets were downloaded and quality control metrics were assessed using Cell Ranger and FastQC, followed by integration with MultiQC. Reads were pre-processed using kallisto-bustools to align to the macaque genome, quantify mapped reads, and generate the cellxgene expression matrix. Cell quality control and downstream analyses were performed using the SCANPY suite, except as noted. To retain smaller resting cells with lower sequencing depth, cells were filtered at minimum cutoffs of 500 unique molecular identifier (UMI) counts and 300 genes per cell. Doublet removal was performed using Scrublet. Cells that passed quality control were merged for downstream analysis. Downstream single-cell RNA-seq analysis was performed using the Python-based SCANPY and AnnData toolkits^91,97,98^

UMAP projection was performed following log1p normalization, calculation of Pearson residuals, identification of highly variable genes, mean-centering and scaling, regression of percent mitochondrial counts and cell cycle scores, dimensionality reduction using PCA, and batch effect removal using BBKNN and Harmony, both of which reported similar results^99,100^. Clustering was performed using the Leiden algorithm. Marker genes were identified using SCANPY’s rank_genes_groups function with default settings, and p-values were adjusted using the Benjamini-Hochberg method^98^. Clusters were annotated based on cell type markers reported in the literature. Pathway analysis was conducted using the gseapy tool and mSigDB, KEGG, and Wikipathways gene list databases94,95,101.

To further address the high dropout rate in scRNA-seq studies and the challenges associated with detecting regulatory factors expressed at low levels, composite gene-set scoring was used to quantify the collective expression of senescence and senescence-related gene sets and complement our direct co-expression detection approach. For each cell, we calculated a gene-set score by summarizing the expression levels of all genes within the set using the score_genes function in SCANPY.

RNA trajectory analysis was conducted using the PAGA package in SCANPY. We excluded a subcluster of bystander cells with small number of counts per cell, which likely represented inefficient sequencing reactions, to prevent them from introducing artifactual patterns in our trajectory study (Supplemental Fig. 14A).

#### Selection of SASP genes for use in identification of senescent cells

To ensure an unbiased selection process, we used the list of SASP genes which were obtained from Gorgoulis et al. (2019). Genes belonging to the four classes of inflammatory factors (interleukins, Chemokines, Other inflammatory molecules, and growth factors and regulators) were selected. Four additional genes, which were more recently shown to be associated with SASP, were added to the list (CCL5, CD163, SPP1, CSF1). Among the genes in this extended list, those represented in the single cell RNA-seq dataset were identified (Fig. S15). Genes which were expressed in more than 20 cells and showed at least 1.3 fold difference in expression between saline-treated and SIVE samples were chosen for use in identification of senescent cells.

#### Selection of control gene sets for senescence genes

To generate an appropriate background control gene set, we matched each gene from our target list of to a control gene based on expression abundance using a custom computational script. First, the mean expression across all cells was calculated for every gene detected in the dataset. For each target gene, a pool of potential control genes was identified, excluding the target genes themselves and any genes previously selected as controls. This pool was filtered to retain only those genes whose mean expression level was within a defined relative tolerance (±2.5%) of the respective target gene’s mean expression. From the qualifying candidates for each target gene, one control gene was selected randomly, ensuring that each control gene was unique within the final matched set. This procedure yielded a control gene set closely matched to the target genes in terms of overall expression abundance, thereby controlling for potential expression level-dependent effects in subsequent analyses. We generated four sets of random genes using this approach, which shared a maximum of 1 gene between each gene list pair.

#### Differential expression tests between saline-treated and SIV+ and SIV-SIVE cells

SIV+ samples were selected as microglial cells with detectable expression of HIV genes. The rest of the microglial cells in the SIVE sample were designated as bystander cells. Each of the saline-treated, SIV+ and bystander cells were converted to pseudobulk, followed by differential expression test using DEseq2. The results were filtered using an FDR cutoff of 0.05 and differential expression level of 1.5 and 2 fold in parallel tests. Neither threshold yielded any significant differentially expressed gene between SIV+ and bystander cells.

#### Statistical modeling

To address the challenges of dropout and technical variability inherent in single-cell RNA sequencing data, we implemented a custom hurdle model that integrates two generalized linear mixed models (GLMMs). In the first step, we modeled the binary outcome of gene detection by classifying each cell as either “positive” or “negative” for the gene of interest. We fitted a binomial GLMM in which the response variable (positive/negative) was modeled as a function of the experimental condition (e.g., SIV vs. saline) and the total transcript count per cell (a proxy for library size), with a random intercept for each donor to account for inter-individual variability. This approach allowed us to adjust for differences in detection sensitivity across cells while quantifying the proportion of cells expressing the marker.

For cells identified as positive, we implemented a second GLMM to model the continuous expression levels. Depending on the distribution of the expression data, an appropriate model (e.g., using a negative binomial distribution) was applied, again incorporating total transcript counts as a covariate and donor as a random effect. The two components of the hurdle model were then integrated—using a likelihood-based framework—to provide an overall differential expression assessment that captures both changes in the detection rate and differences in expression intensity among positive cells. This integrated approach enables a comprehensive evaluation of gene expression dynamics by considering both the proportion of expressing cells and the magnitude of expression, thereby providing a more robust analysis of differential expression between conditions.

A Pearson chi-square test of independence was conducted to examine differences in the proportions of senescent cells among the study groups. Kolmogorov–Smirnov test was used to compare the co-expression distributions between different experimental groups.

For Investigating the enrichment pattern for SIV+ cells in those with senescence signature, a 2×2 contingency table was constructed using counts of cells classified by SIV status (SIV+ versus SIV–) and senescence phenotype (senescence-positive versus senescence-negative). Fisher’s exact test was then applied to assess the association between these categorical variables. This test was implemented using the fisher_exact function from the scipy.stats module in Python, which computes an exact odds ratio and two-tailed p-value. This approach was chosen because it is well-suited for categorical data analysis when some cells have low counts, providing a robust measure of association without relying on large-sample approximations.

