## supplemental-figures for "HIV infection in microglia leads to senescence, triggering activation of neurotoxicity pathways"

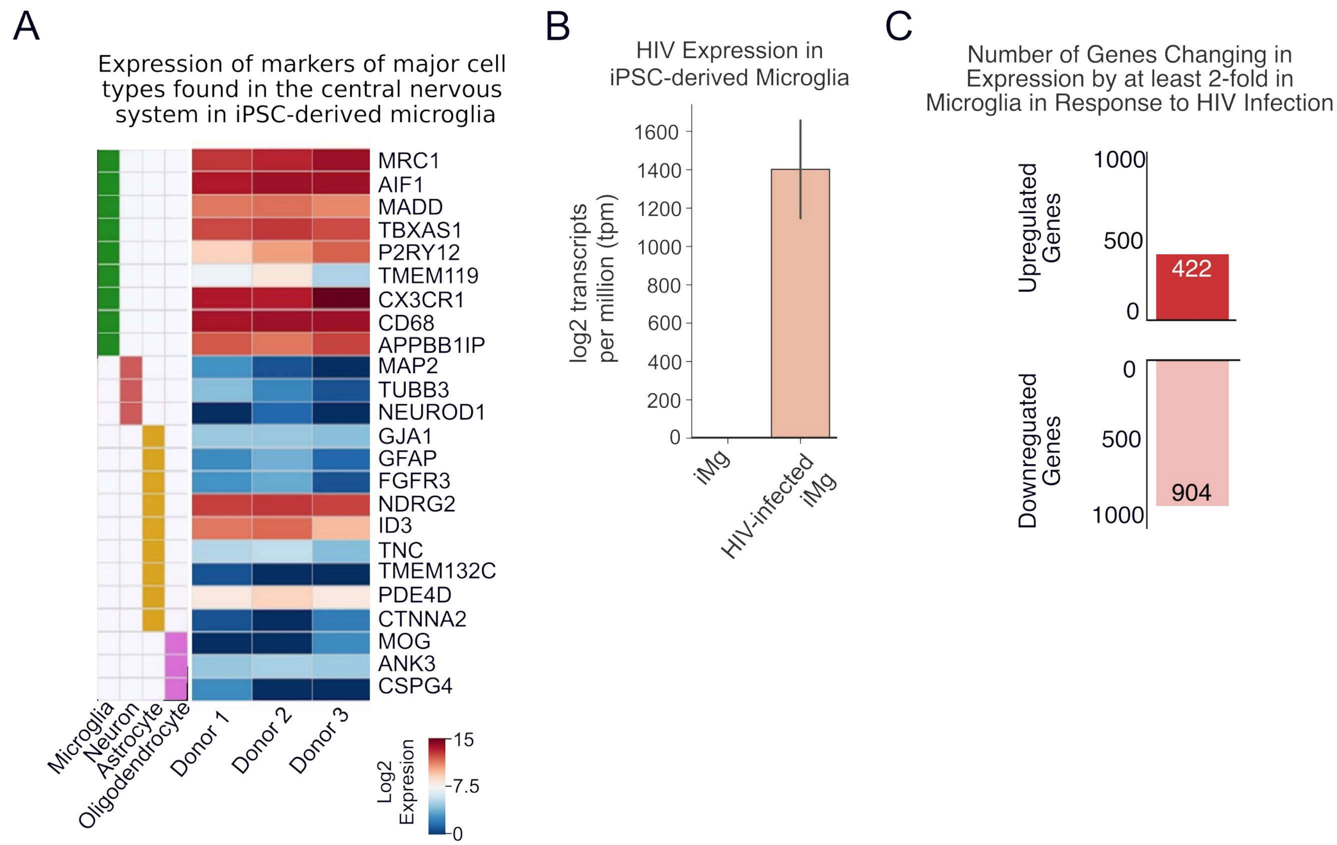

**Supplemental Figure 1.** Validation of Microglial Cell Types and impact of HIV infection. (A) Heatmap showing expression of microglia, neuron, astrocyte, and oligodendrocyte genes in uninfected iPSC-derived microglia. The heatmap on the right side of the figure shows the expression level of the genes shown to the right of the panel. The annotation matrix to the left of the heatmap identifies the cell types (listed at the bottom) corresponding to each group of markers. (B) Barplot showing expression level of HIV in mock infected (left) and HIV-infected (right) samples. The error bar represents standard deviation among the three donors. (C) Barplot showing the number of upregulated and downregulated genes in HIV infected sample compared to mock infected control.

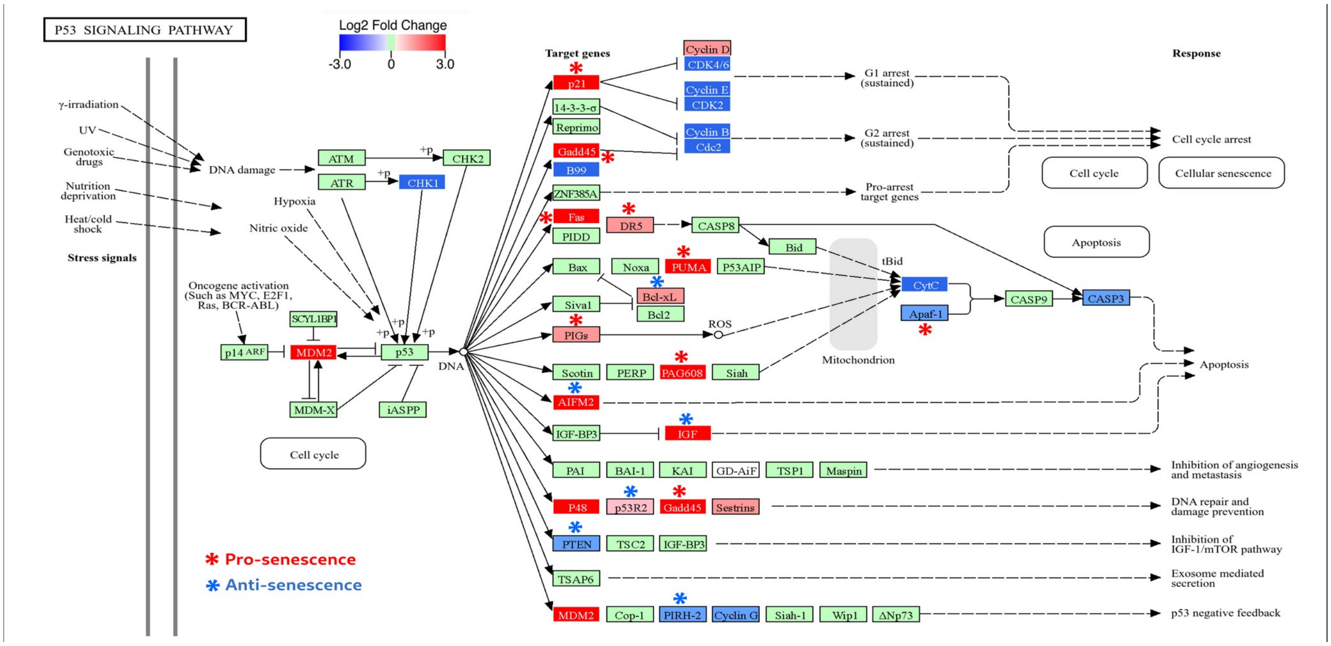

**Supplemental Figure 2.** p53 signaling cascade with the changes in gene expression after HIV infection superimposed on each node in the plot. Green, red and blue colored nodes represent genes showing no change, up- and downregulation after HIV infection compared to mock infected, respectively. Red and blue asterisks mark pro- and anti-senescence genes, respectively. The p53-induced pathways that lead to cell cycle arrest and cellular senescence, concomitant with downregulation of caspase3, are shown. Color bar at the top shows the direction and magnitude of the changes in gene expression for each node. The pathway graph is obtained from KEGG network database.

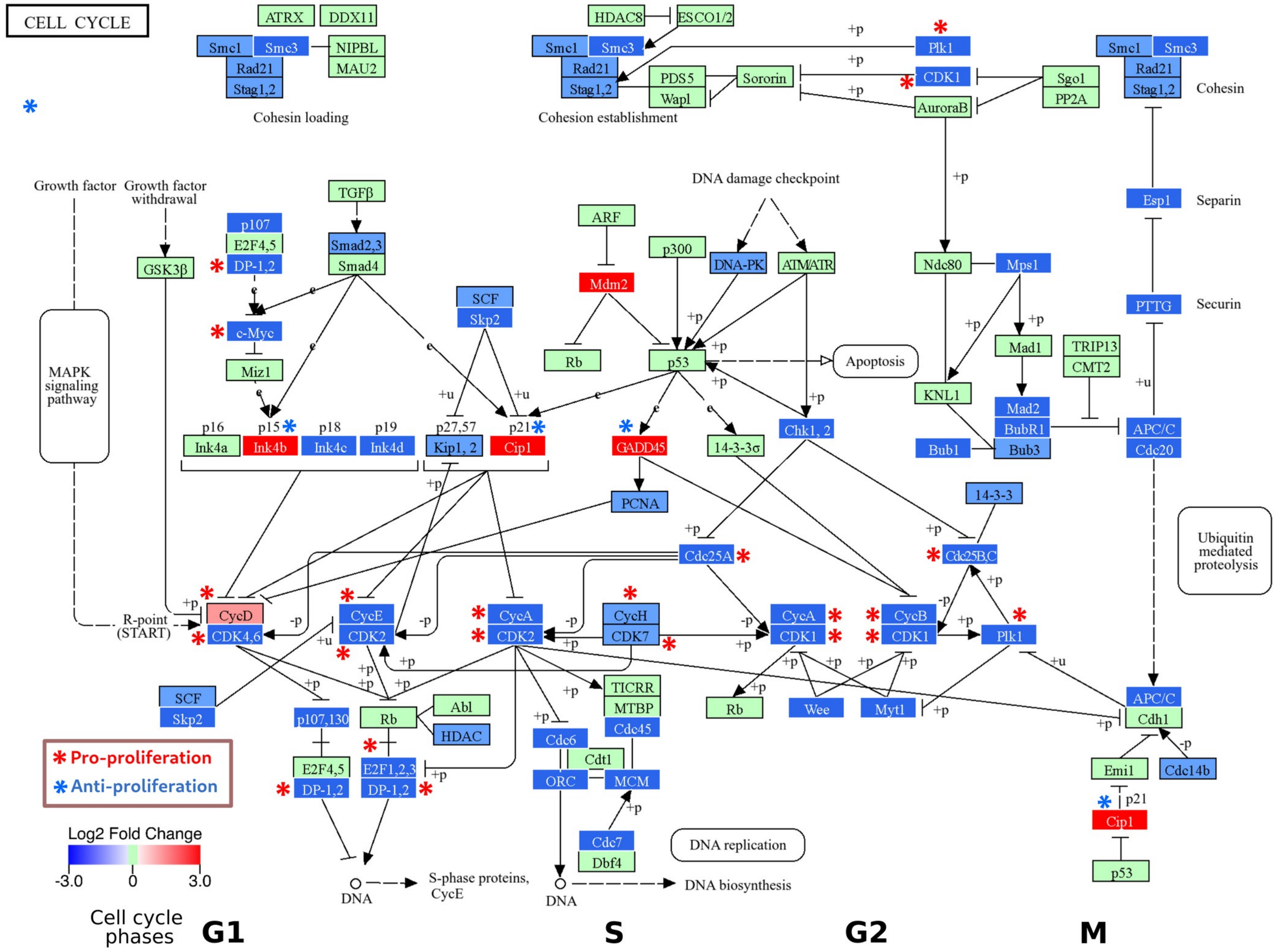

**Supplemental Figure 3**. Cell cycle regulatory pathways with the changes in gene expression after HIV infection superimposed on each node in the plot, showing the marked downregulation of positive regulators of proliferation. These include cyclins and cyclin-dependent kinases, along with c-MYC, DP-1 and 2, PIK1 and CDD25A, B and C (marked by red asterisks). Upregulated genes include Cip1/p21/CDKN1A, GADD45 and p15/Ink4a/CDKN2B, three major inhibitors of cell cycle and markers of cellular senescence (marked by blue asterisks). Color bar at the bottom left shows the direction and magnitude of the changes in gene expression for each node. Green, red and blue colored nodes correspond to genes showing no change, up- and downregulation after HIV infection compared to mock infected, respectively. The cellular events are organized to represent G1, S, G2 and M phases of cell cycle from left to right, as marked at the bottom of the plot. The pathway graph is obtained from KEGG network database.

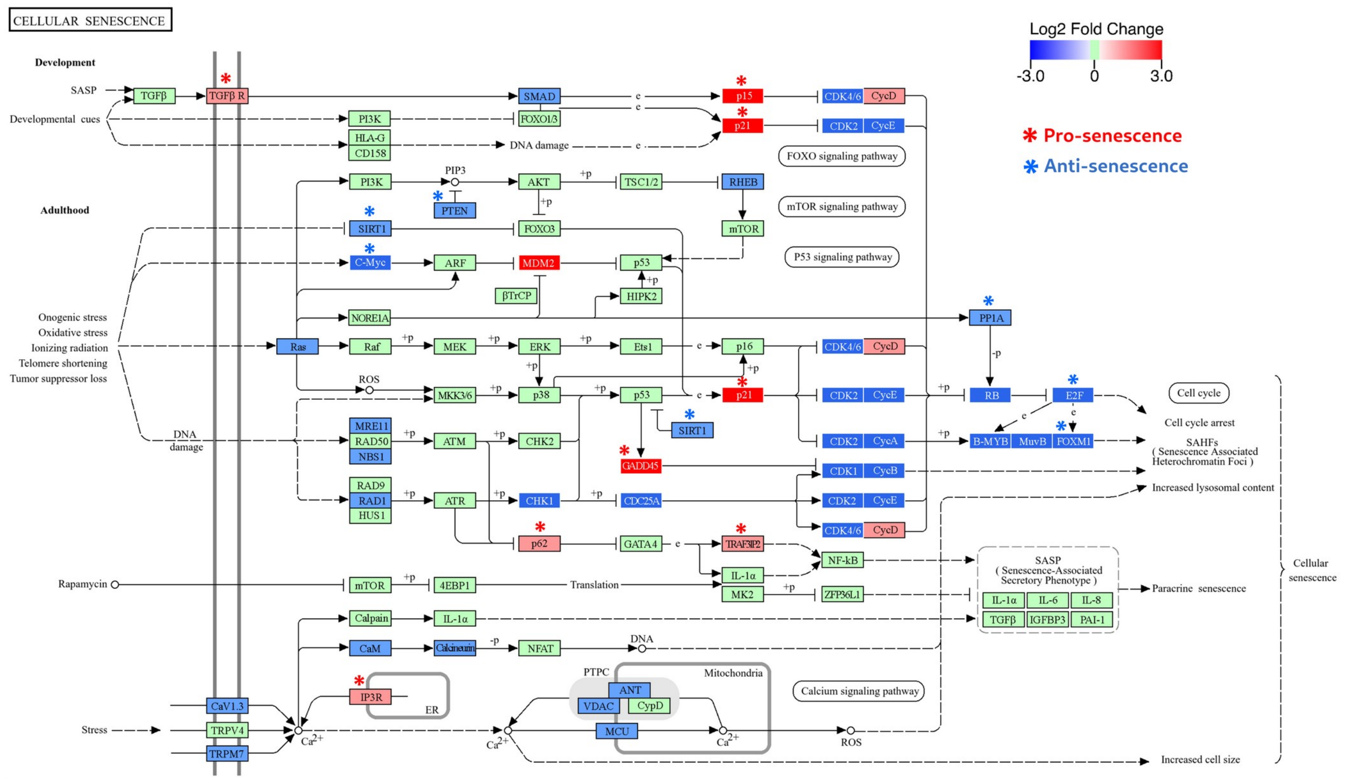

**Supplemental Figure 4.** Pathways and factors regulating cellular senescence, with the changes in gene expression after HIV infection superimposed on each node in the plot. Green, red and blue colored nodes correspond to genes showing no change, up- and downregulation after HIV infection compared to mock infected, respectively. Red asterisks mark the pro-senescence factors including p21/CDKN1A, p15/CDKN2B, GADD45, p62/SQSTM1, TGFb receptor, IP3R and TRAF3IP2, all of which are upregulated. On the other hand, key negative regulators of senescence such as SIRT1, c-MYC, PP1A, E2F family, FOXM1 and PTEN are strongly downregulated after HIV infection (marked by blue asterisks). Color bar at the top right shows the direction and magnitude of the changes in gene expression for each node. The pathway graph is obtained from KEGG network database.

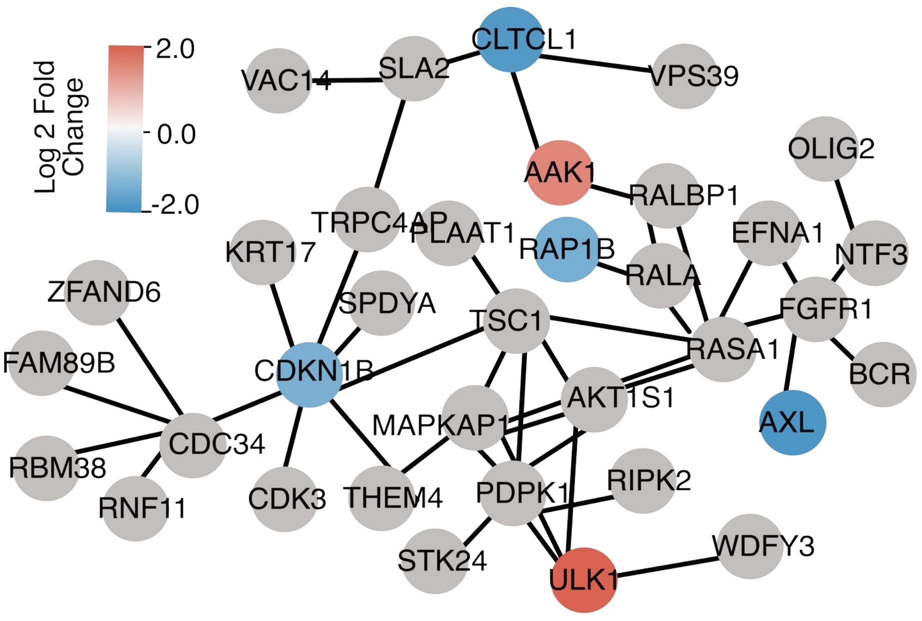

**Supplemental Figure 5.** MTOR pathway is not downregulated in HIV-infected iPSC-derived microglial cells. Each node represents one of the genes involved in the mTOR pathway. The edges (thin lines) connecting the different nodes represent co-expression or direct interaction. Red and blue colors represent up and down-regulated genes, while gray nodes showed no changes in expression.

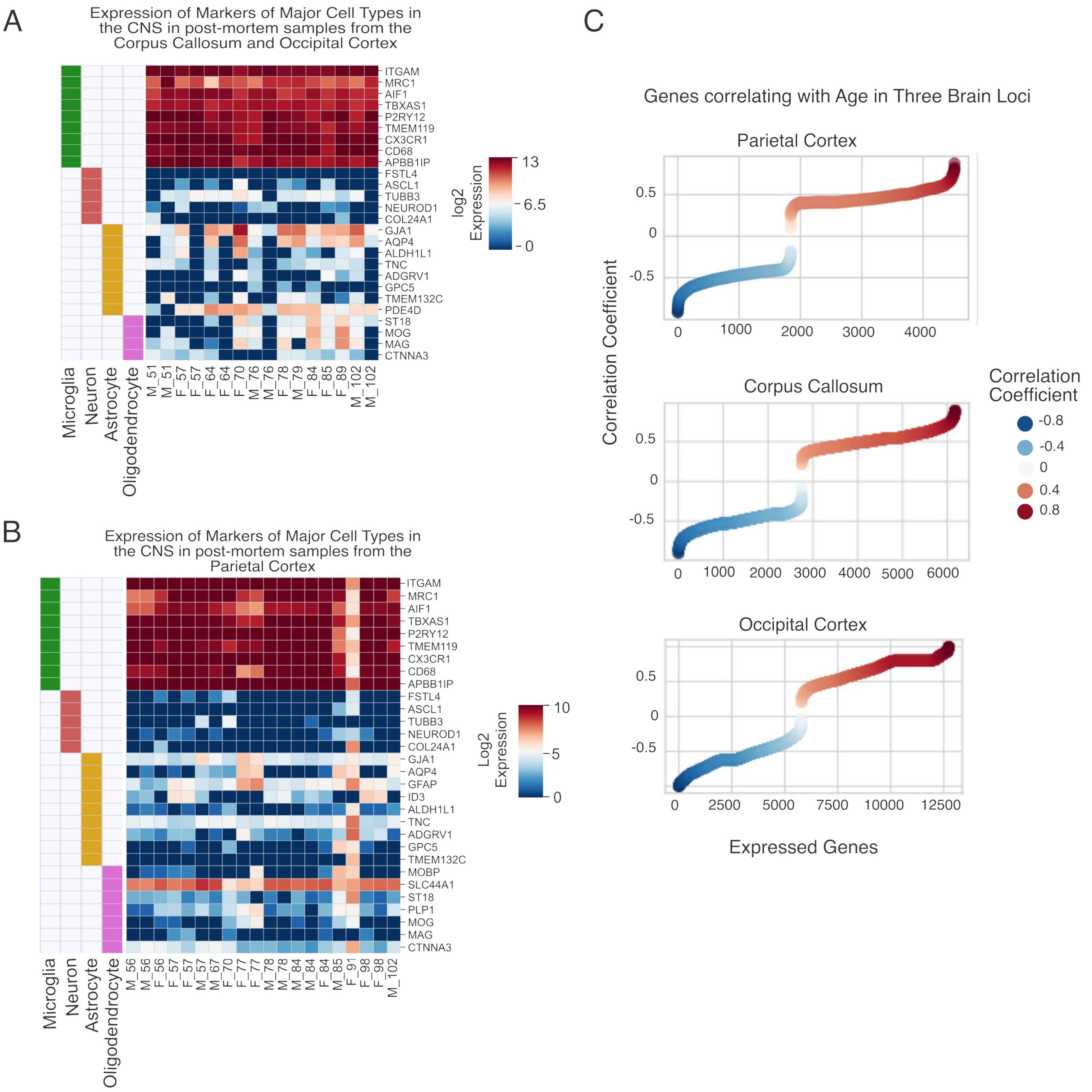

**Supplemental Figure 6.** Validation of microglia in aging datasets and identification of aging-correlated genes. A,B. Heatmaps showing expression of microglia, neuron, astrocyte, and oligodendrocyte marker genes in the two aging datasets. The annotation plot to the left of the heatmap indicates the cell type specificity of each gene. Male and female donors are marked as M and F, respectively. The numbers in the X axis legend are biological age of each donor. (C) S-Curves showing the number of genes per brain locus that positively or negatively correlate with age (red and blue lines, respectively).

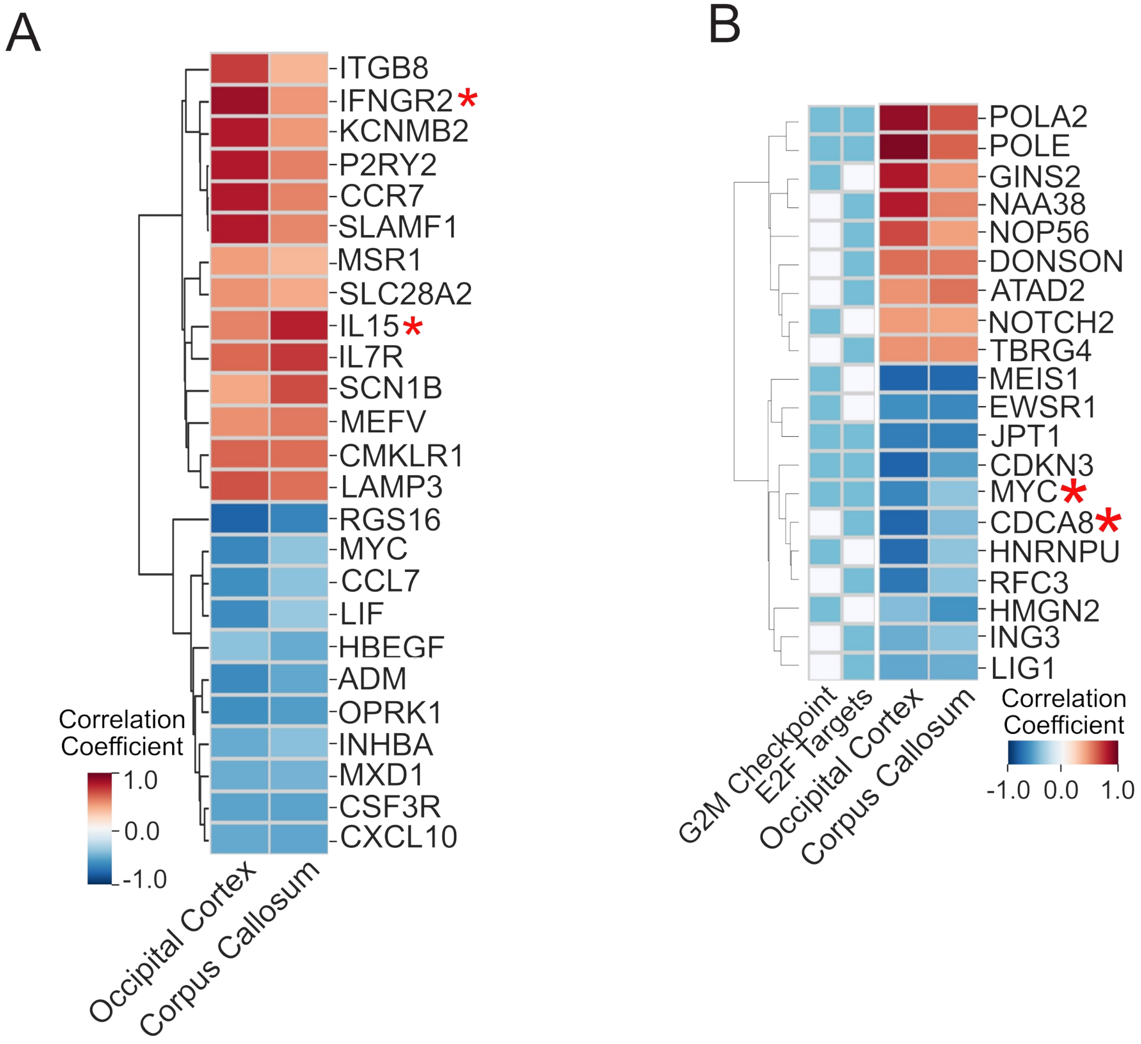

**Supplemental Figure 7.** Identification of expression pattern of Aging-associated inflammatory and proliferative genes in primary human microglia obtained from post-mortem tissues.

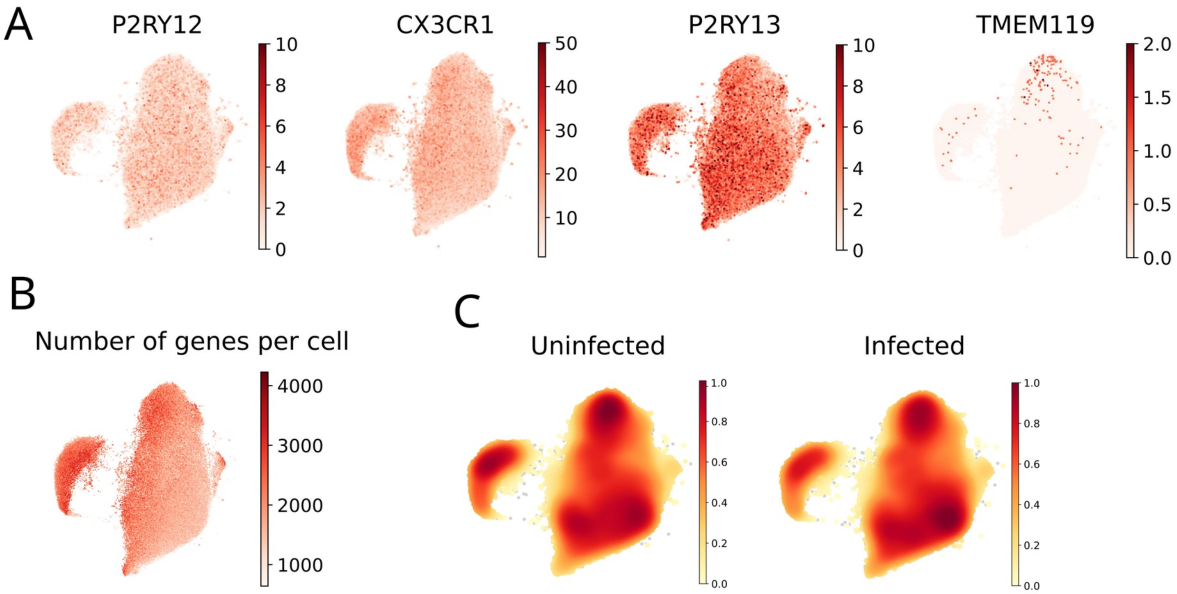

**Supplemental Figure 8. Validation of iPSC-derived microglia.** A. Microglial markers are robustly expressed in iPSC-derived microglial cells. B. Number of genes per cell indicates a range of sequencing depths in cells, necessitating the inclusion of transcriptomic depth as a factor in statistical studies. C. Density plots indicate the similar distribution of cells across the clusters in cells from the two experimental conditions.

Co-expression values for CDKN1A and CDKN2A per cluster and in aggregate

| | CDKN1A + CDKN2B double positive | Total cells per group | % double positive per group |
|---|---|---|---|
| **In all clusters** | | | |
| **HIV+ (total)** | 131 | 3320 | 3.95 |
| **Bystander (total)** | 1796 | 41040 | 4.38 |
| **Uninfected (total)** | 1061 | 47336 | 2.24 |
| **Per cluster** | | | |
| **HIV_0** | 20 | 1448 | 1.38 |
| **HIV_1** | 50 | 1509 | 3.31 |
| **HIV_2** | 61 | 363 | 16.80 |
| **Bystander_0** | 439 | 23312 | 1.88 |
| **Bystander_1** | 337 | 12187 | 2.77 |
| **Bystander_2** | 1020 | 5541 | 18.41 |
| **Uninfected_0** | 212 | 25068 | 0.85 |
| **Uninfected_1** | 213 | 15040 | 1.42 |
| **Uninfected_2** | 636 | 7228 | 8.80 |
| **Total** | 2988 | 91696 | 3.26 |

**Supplementary Figure 9.** Co-expression pattern of CDKN1A/p21 and CDKN2A/p16 in each experimental group and cluster.

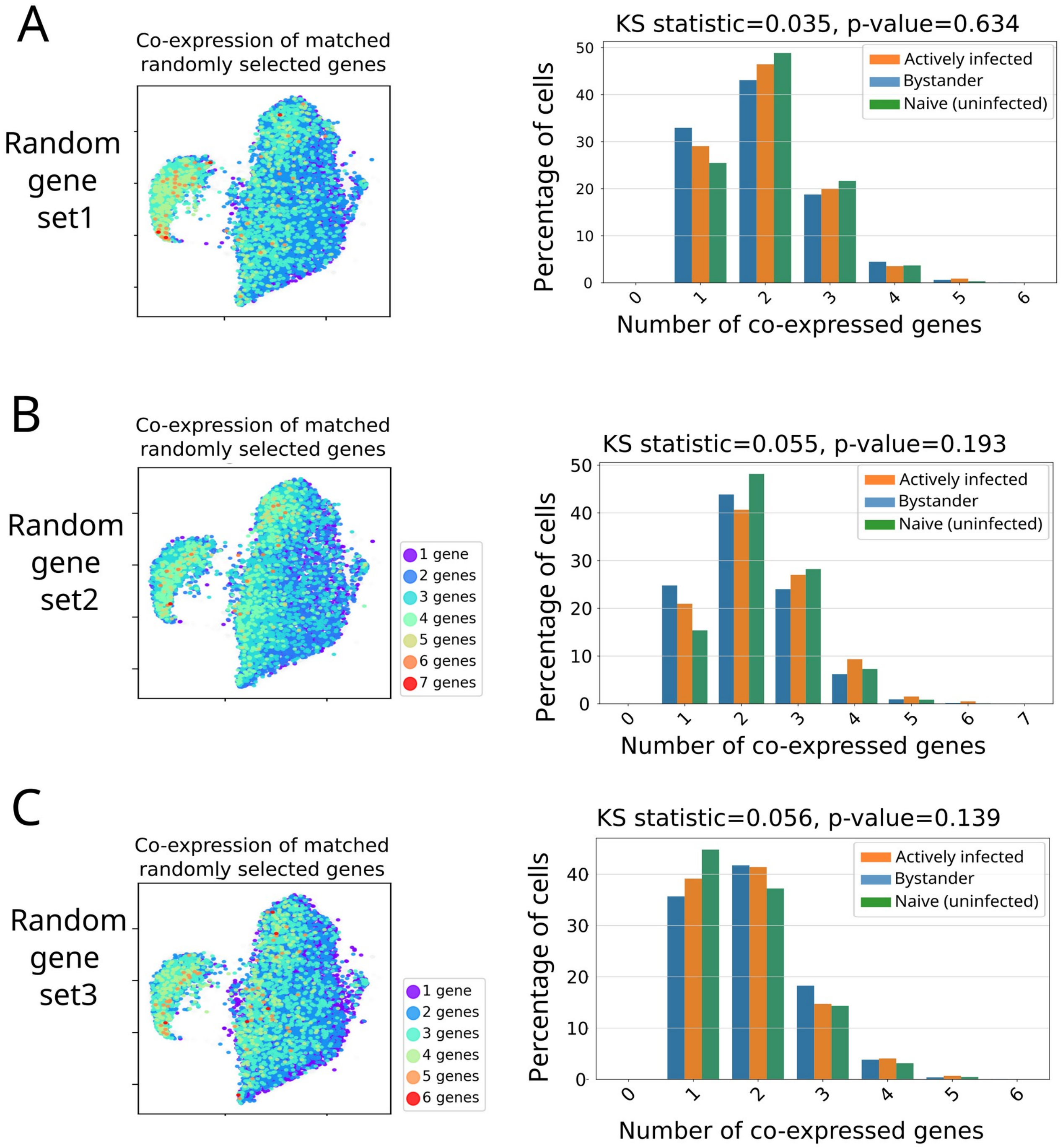

**Supplementary Figure 10.** Co-expression pattern of control gene set show distributions distinct from those obtained with senescent cells. Kolmogorov-Smirnov (KS) test values for comparison of distribution of actively infected and naive cells are shown for each set. The control genes were selected by first grouping genes with expression patterns similar to each of our 11 senescence genes, then randomly selecting one representative from each group (see Methods).

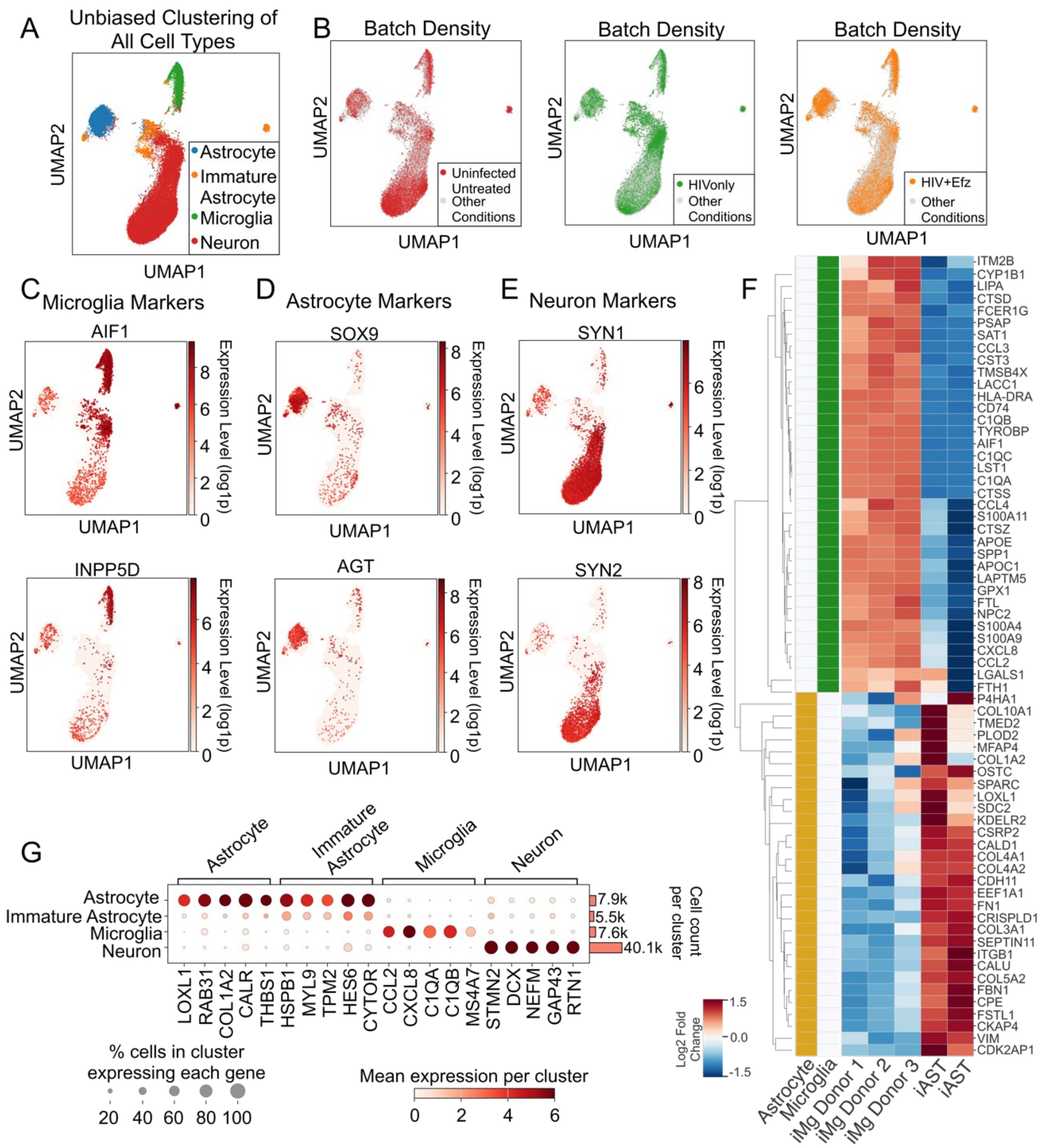

**Supplemental Figure 11**. Cell type assignment in single cell RNA-sequencing data of HIV-infected iPSC-derived human microglia. (A) UMAP of unbiased clustering of all cell types. (B) UMAPs showing batch densities of different treatments. (C) UMAP of expression of microglia markers AIF1 and INPP5D. (D) UMAP of expression of astrocyte markers SOX9 and AGT. (E) UMAP of neuron markers SYN1 and SYN2. (F) Heatmap showing the bulk RNA-seq expression of the top 25 genes expressed by microglia and top 25 genes expressed by astrocytes identified in single cell RNA-seq analyses. (G) Dotplot showing expression of cellular markers for each cell type.

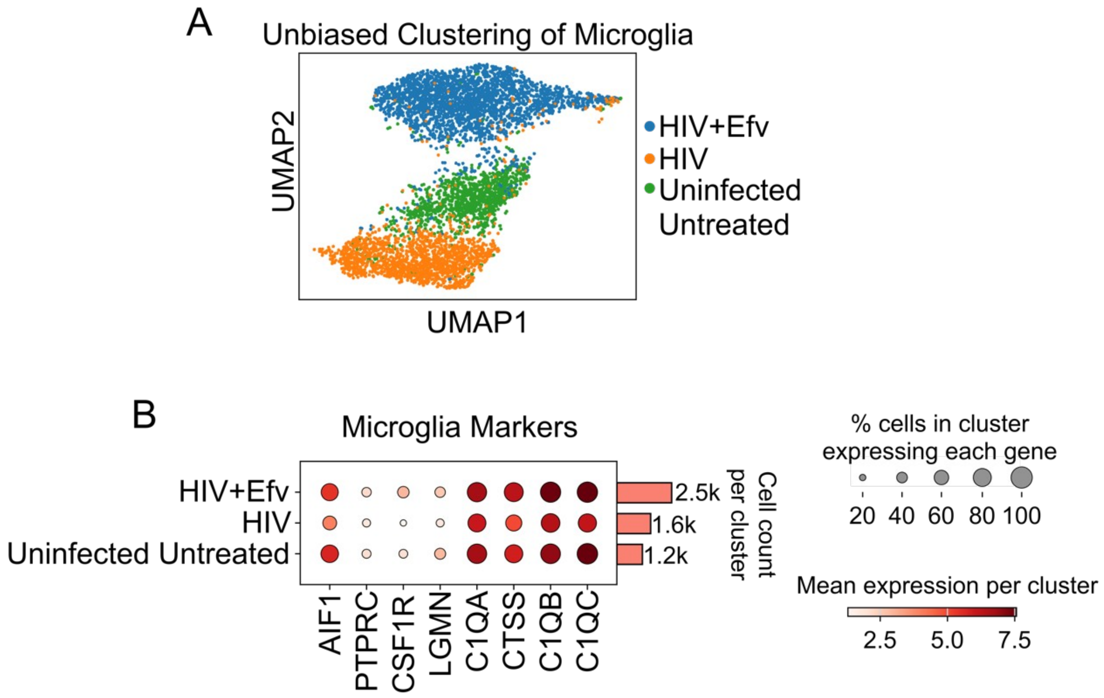

**Supplemental Figure 12.** Microglia clustering and marker expression. (A) UMAP of microglia by treatment. (B) Dotplot of expression of microglia marker genes in each treatment.

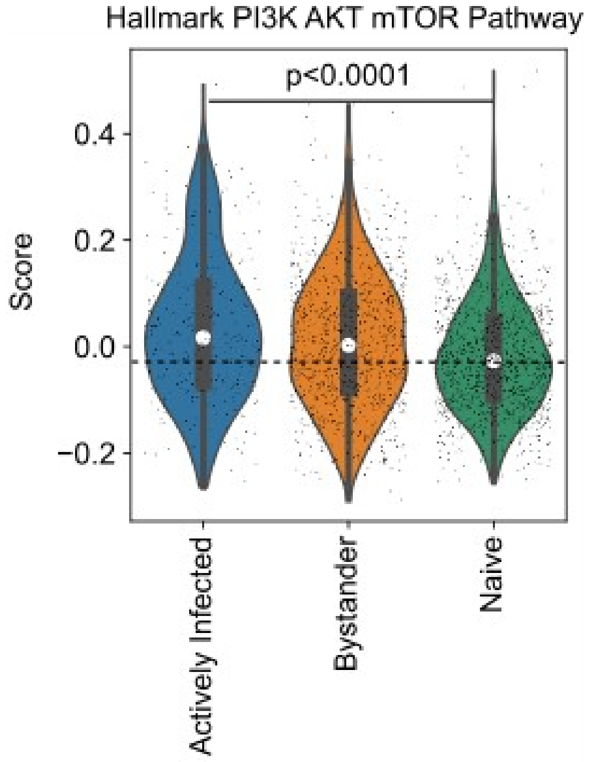

**Supplemental Figure 13.** Aggregate score of expression of genes in the Hallmark PI3K AKT mTOR pathway in HIV-infected iPSC-derived microglia.

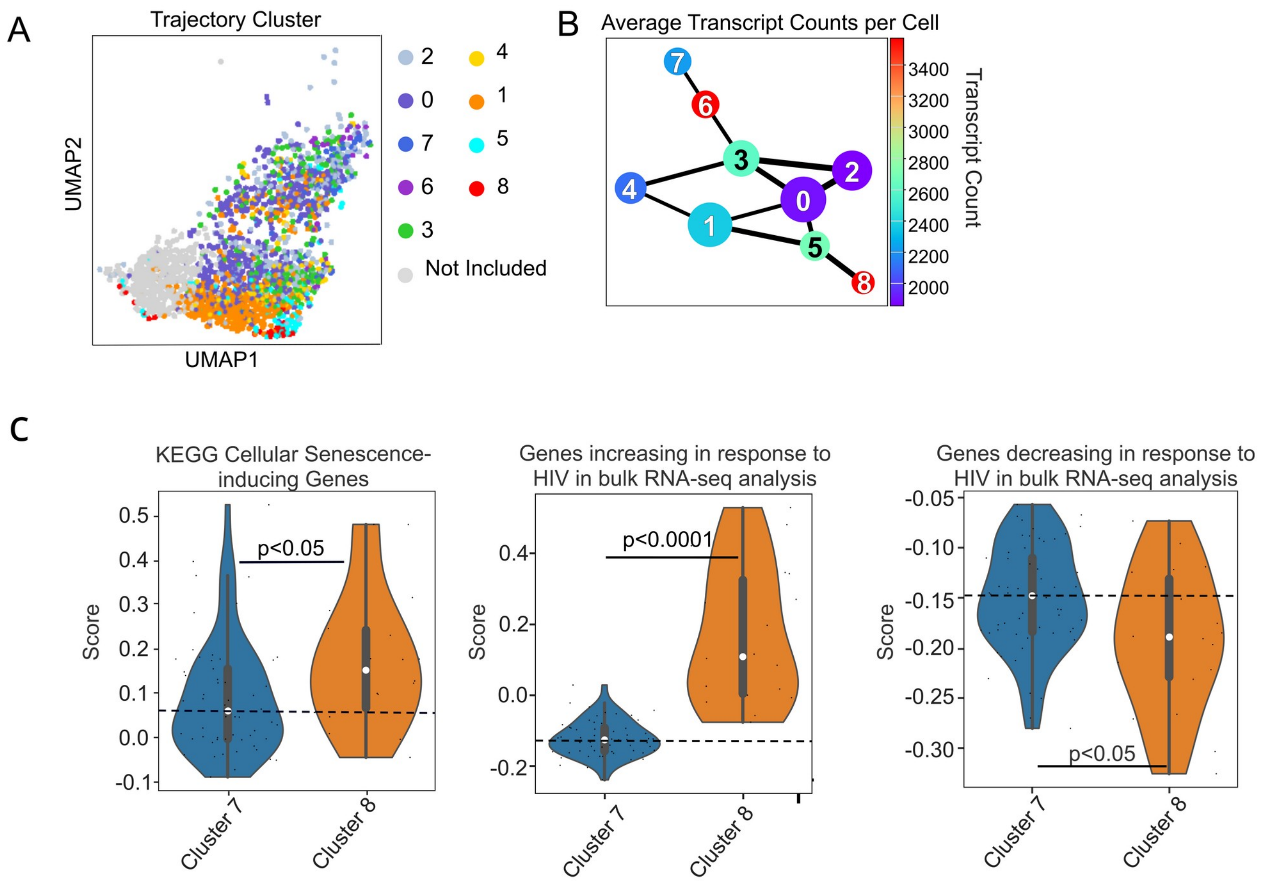

**Supplemental Figure 14.** A. UMAP plot showing trajectory clusters. A subcluster (shown in gray) which contained cells with low sequencing depth relative to other clusters was omitted from the study. B. Trajectory analysis showing average transcript counts per cell. C. Violin plot showing expression of KEGG cellular senescence-inducing genes, genes increasing in response to HIV identified in the bulk RNA-seq analysis, and genes decreasing in response to HIV identified in the bulk RNA-seq analysis in the first and last clusters of the trajectory.

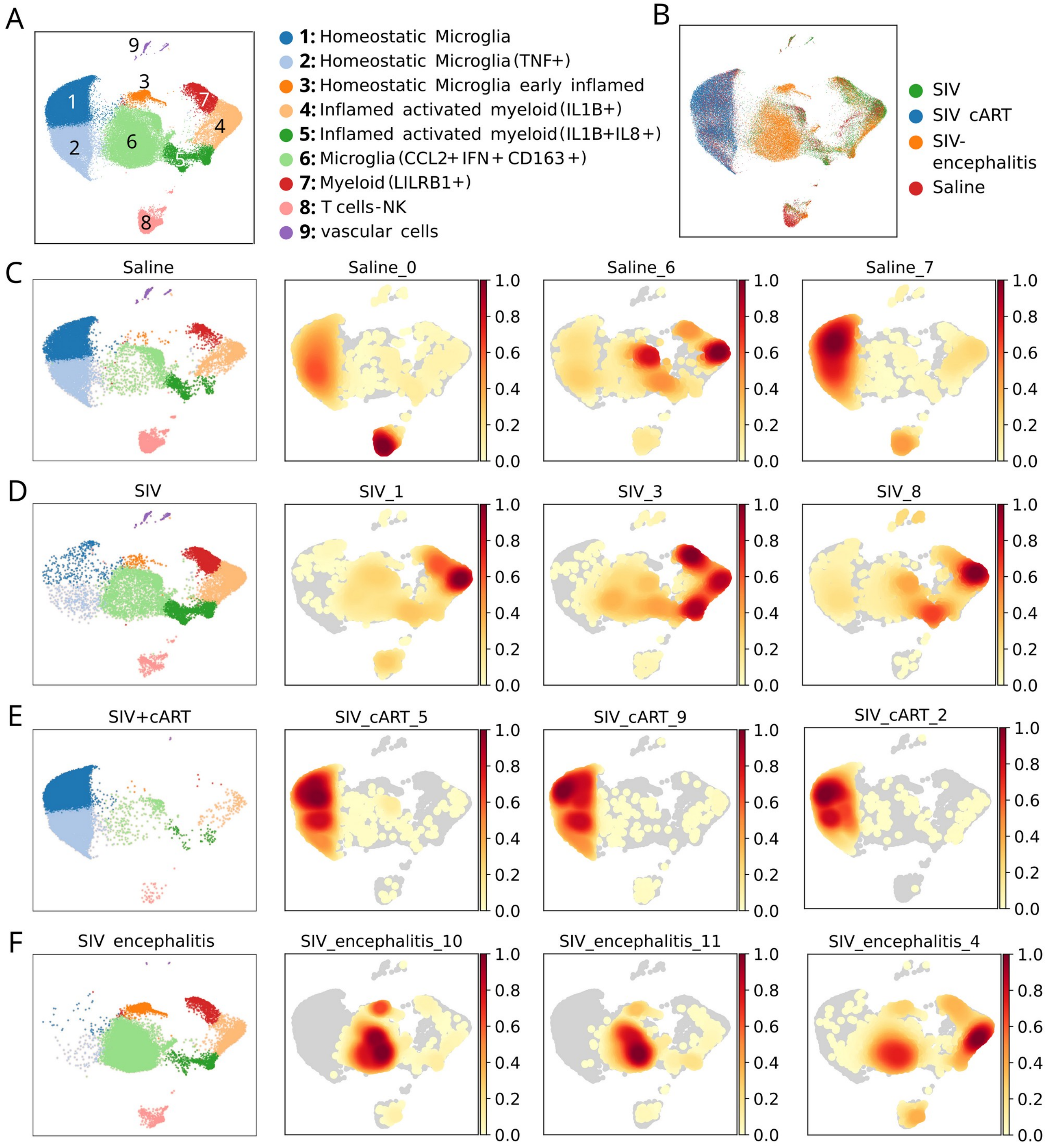

**Supplementary Fig. 15.** Clustering of macaque samples. A. Identification of clusters and cell type assignment. B. UMAP showing the position of each treatment group. C-F, UMAP showing the position of cells belonging to each treatment group, along with UMAPs for each individual macaque study subject.

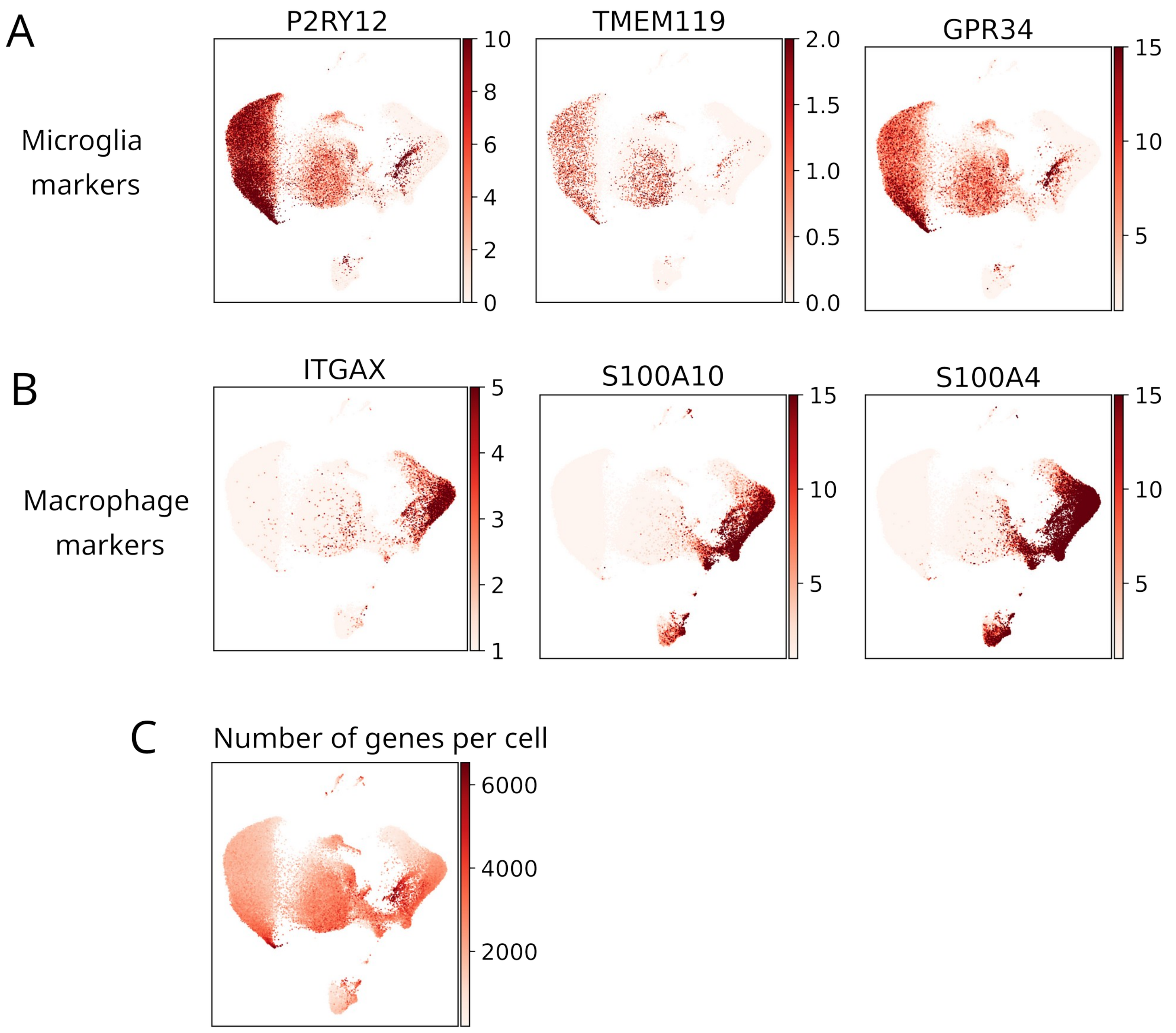

**Supplemental Figure 16.** A and B. Lineage marker genes show the microglial clusters and clusters which likely contain non-microglial myeloid cells. C. UMAP showing the number of genes represented in each cell.

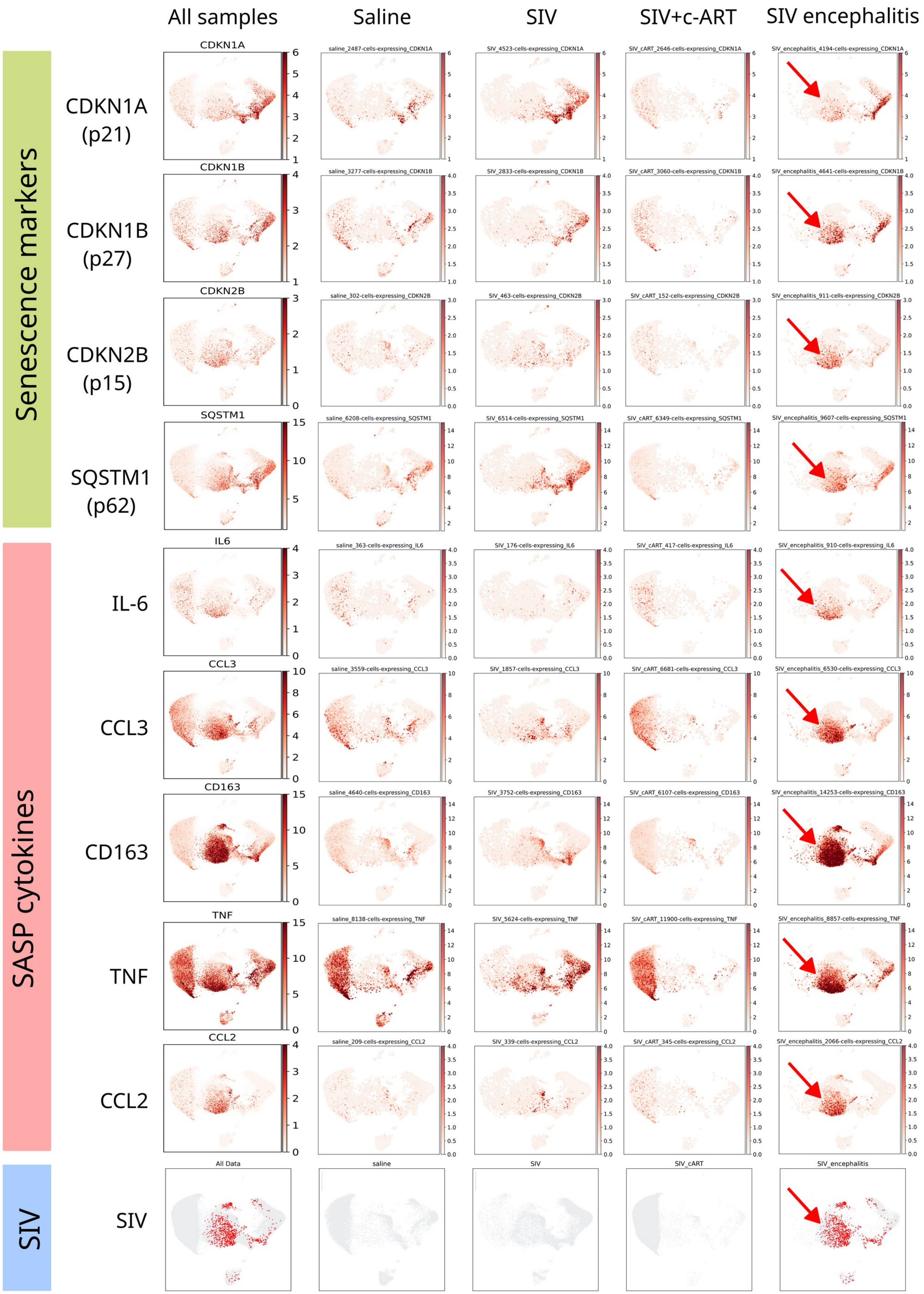

All samples
Saline
SIV
SIV+c-ART
SIV encephalitis
Senescence markers
CDKN1A (p21)
CDKN1B (p27)
CDKN2B (p15)
SQSTM1 (p62)
SASP cytokines
IL-6
CCL3
CD163
TNF
CCL2
SIV
CDKN1A
saline_2487-cells-expressing_CDKN1A
SIV_4523-cells-expressing_CDKN1A
SIV_cART_2646-cells-expressing_CDKN1A
SIV_encephalitis_4194-cells-expressing_CDKN1A
CDKN1B
saline_3277-cells-expressing_CDKN1B
SIV_2833-cells-expressing_CDKN1B
SIV_cART_3060-cells-expressing_CDKN1B
SIV_encephalitis_4641-cells-expressing_CDKN1B
CDKN2B
saline_302-cells-expressing_CDKN2B
SIV_463-cells-expressing_CDKN2B
SIV_cART_152-cells-expressing_CDKN2B
SIV_encephalitis_911-cells-expressing_CDKN2B
SQSTM1
saline_6208-cells-expressing_SQSTM1
SIV_6514-cells-expressing_SQSTM1
SIV_cART_6349-cells-expressing_SQSTM1
SIV_encephalitis_9607-cells-expressing_SQSTM1
IL6
saline_363-cells-expressing_IL6
SIV_176-cells-expressing_IL6
SIV_cART_417-cells-expressing_IL6
SIV_encephalitis_910-cells-expressing_IL6
CCL3
saline_3559-cells-expressing_CCL3
SIV_1857-cells-expressing_CCL3
SIV_cART_6681-cells-expressing_CCL3
SIV_encephalitis_6530-cells-expressing_CCL3
CD163
saline_4640-cells-expressing_CD163
SIV_3752-cells-expressing_CD163
SIV_cART_6107-cells-expressing_CD163
SIV_encephalitis_14253-cells-expressing_CD163
TNF
saline_8138-cells-expressing_TNF
SIV_5624-cells-expressing_TNF
SIV_cART_11900-cells-expressing_TNF
SIV_encephalitis_8857-cells-expressing_TNF
CCL2
saline_209-cells-expressing_CCL2
SIV_339-cells-expressing_CCL2
SIV_cART_345-cells-expressing_CCL2
SIV_encephalitis_2066-cells-expressing_CCL2
All Data
saline
SIV
SIV_cART
SIV_encephalitis

**Supplementary Figure 17.** Expression pattern of key senescence markers and SASP-associated genes are shown in UMAPs for each treatment condition. The subcluster showing consistent expression of the senescent markers is marked by arrows.

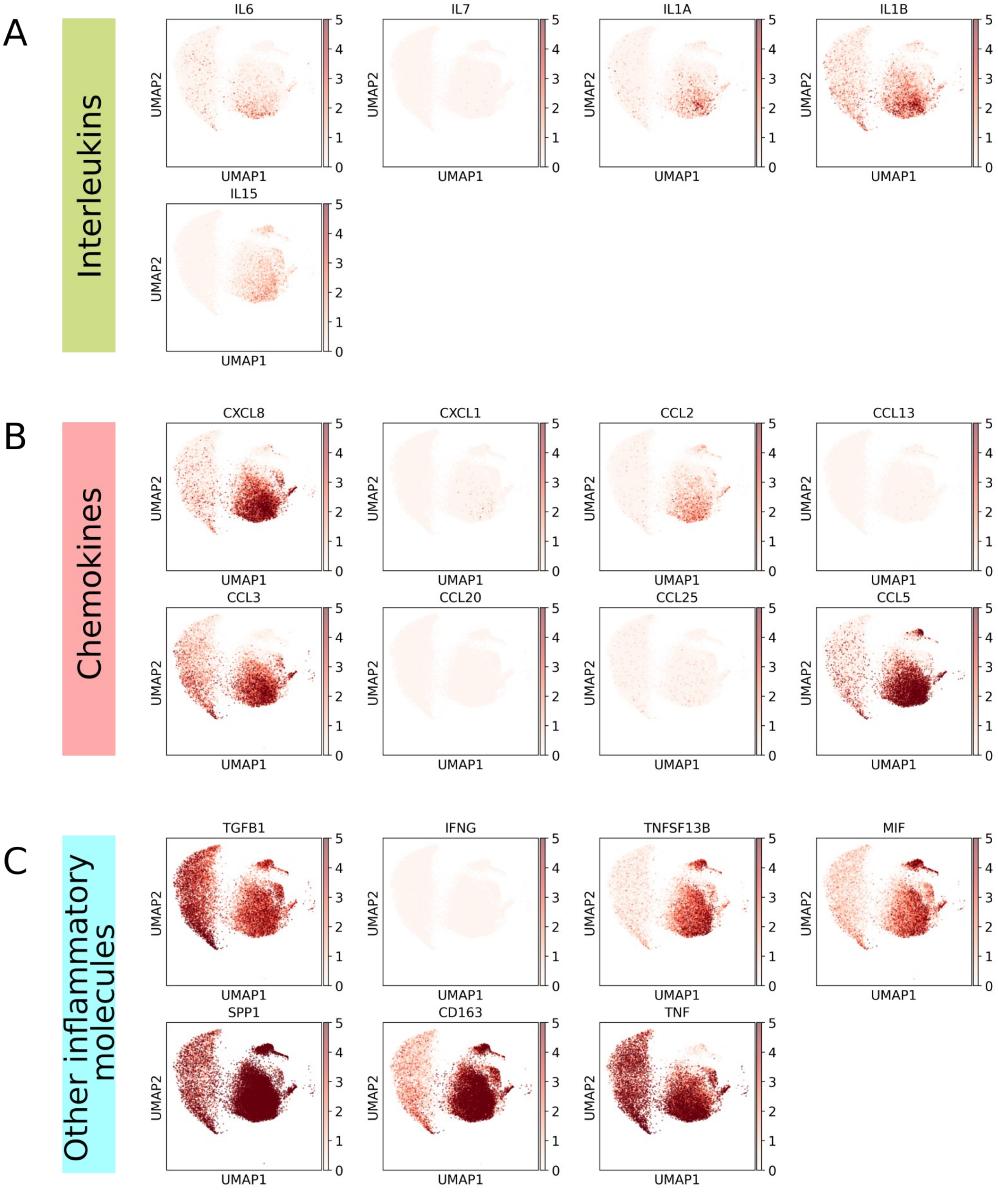
A
Interleukins
IL6
IL7
IL1A
IL1B
IL15
B
Chemokines
CXCL8
CXCL1
CCL2
CCL13
CCL3
CCL20
CCL25
CCL5
C
Other inflammatory molecules
TGFB1
IFNG
TNFSF13B
MIF
SPP1
CD163
TNF
UMAP1
UMAP2
0
1
2
3
4
5

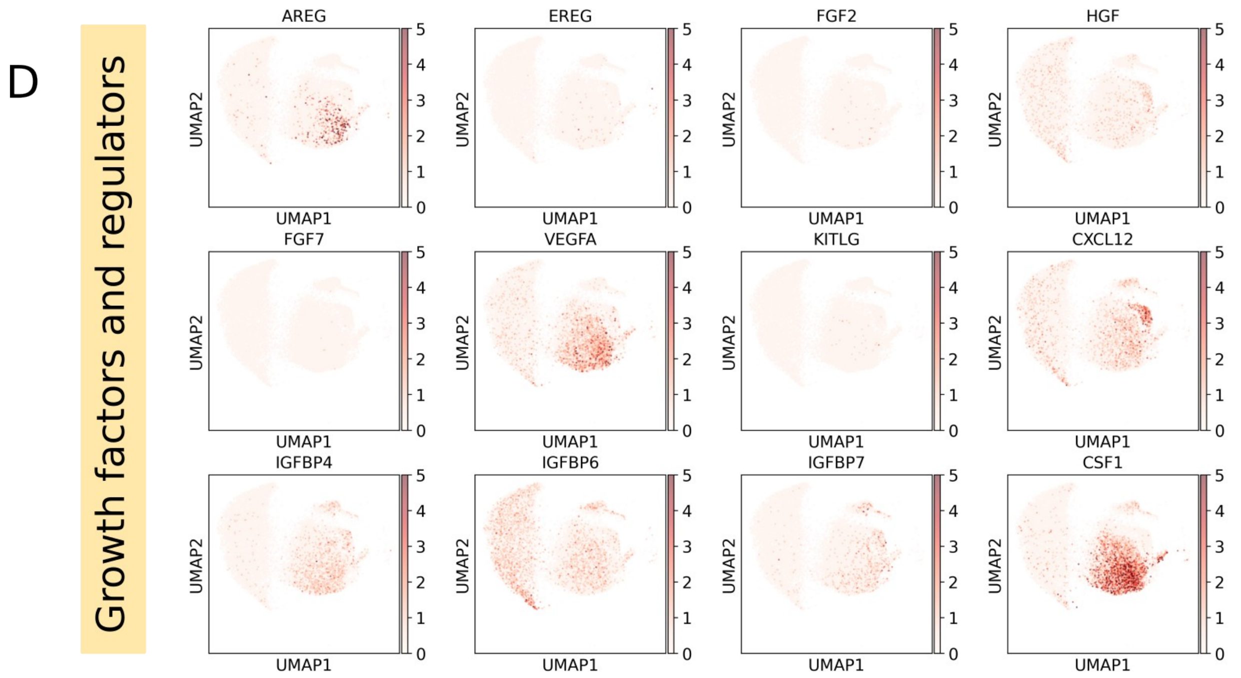

**Fig. S18. The SASP-associated genes used in this study.** The list of SASP genes were obtained from Gorgoulis et al., 2019 and genes represented in the single cell RNA-seq dataset were identified. Genes which were expressed in more than 20 cells and showed at least 1.2 fold difference in expression between saline and SIVE samples were chosen for use in identification of senescent cells (Table 2). A-D: SASP-associated genes belonging to the main four classes of inflammatory molecules (shown using the color bands on the left) per Gorgoulis et al. The color bar to the right of each panel indicates the level of expression in UMI counts.

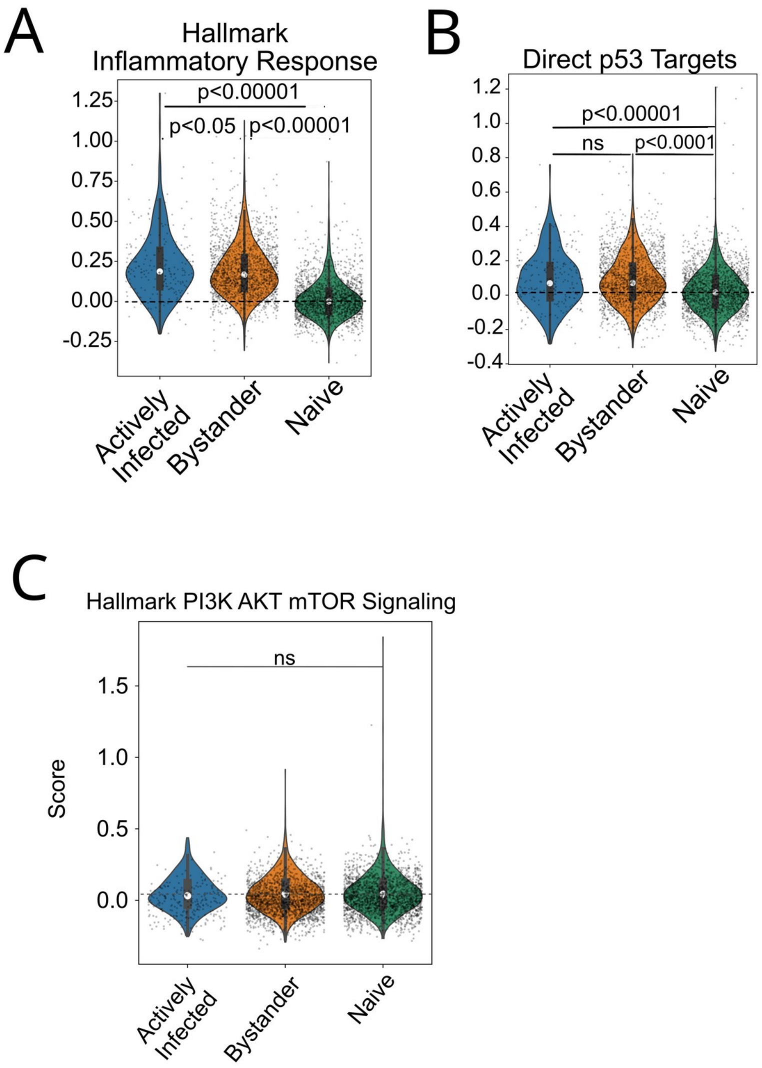

**Supplementary Figure 19. Concomitant elevation of inflammatory response and p53 activity, which play crucial role in induction of senescence, in SIV-infected cells.** Naive = saline-treated cells. All cells in SIV encephalitis (SIVE) sample which did not have detectable expression of SIV genes were assigned to the bystander group. Actively infected = detectable SIV expression. The violin plots indicate the calculated composite expression value for genes in inflammatory response (A), p53 pathway (B) and PI3K-AKT-mTOR signaling (C). p-values for each study were calculated using Mann-Whitney U test. Lack of change in mTOR pathway rules out the possibility of SIV-mediated entry into quiescence. The white dots in violin plots mark the mean value of the group. The small dots represent individual genes. ns = not significant.

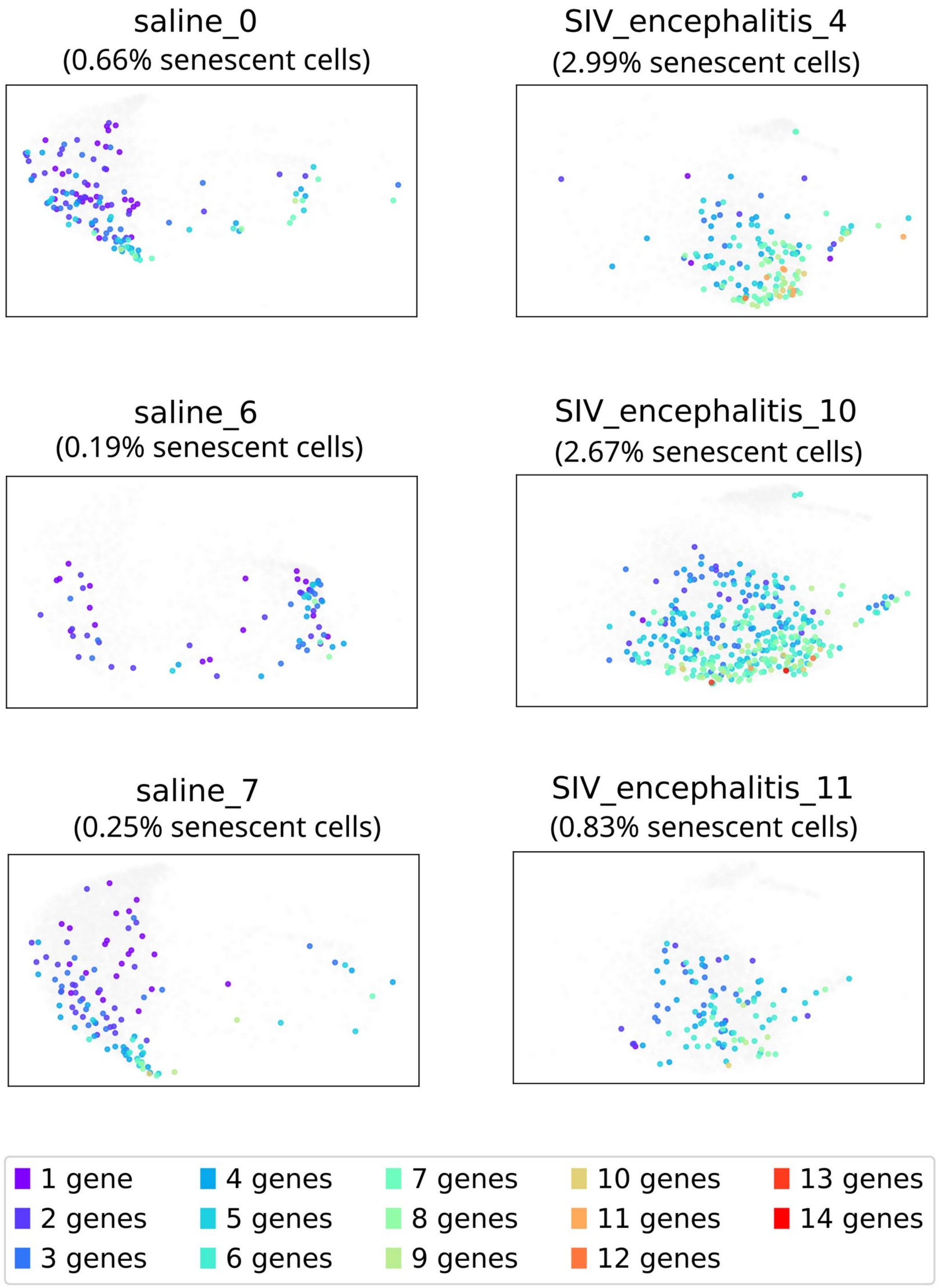

**Supplementary Fig. 20. Comparison of percentage of senescent cells (those with >6 co-expressed senescence-related genes [Table 2], at least one of which must be CDKN1A or CDKN2B) per donor.** The legend at the bottom indicate the color corresponding to each level of co-expression. Only cells which express at least one senescence-related genes are shown in the plots for clarity. The percent senescent cell values are computed as number of microglial cells with >6 co-expressed senescent genes/total number of microglial cells per donor. Statistical analysis of the two groups using Mann-Whitney U test yielded a U statistic of 9 (the largest possible with 3 samples per arm, indicative of complete separation of the two groups) and a p-value of 0.05 (smallest possible for a study with 3 donors per arm).

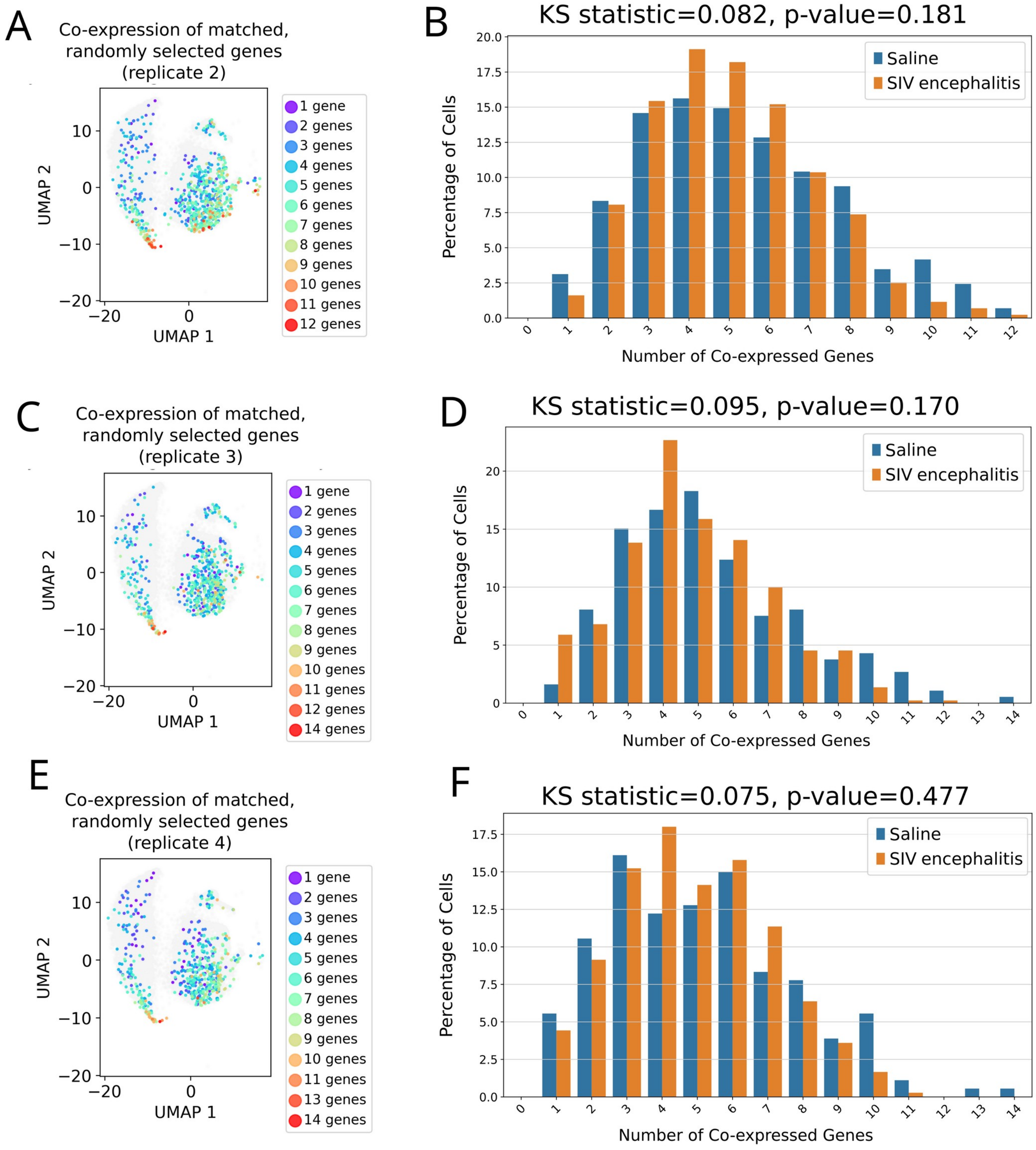

**Supplementary Fig. 21. Multiple sets of randomly selected genes that matched the expression patterns of senescence genes used in Fig. 6E and F show similar coexpression patterns in saline-treated and SIVE samples.** UMAPs and distribution plots are shown for each set of genes. The statistical significance and effect size are indicated above the distribution plots for each sample. Unlike the case with senescence genes, matched random genes show higher co-expression values in saline samples.

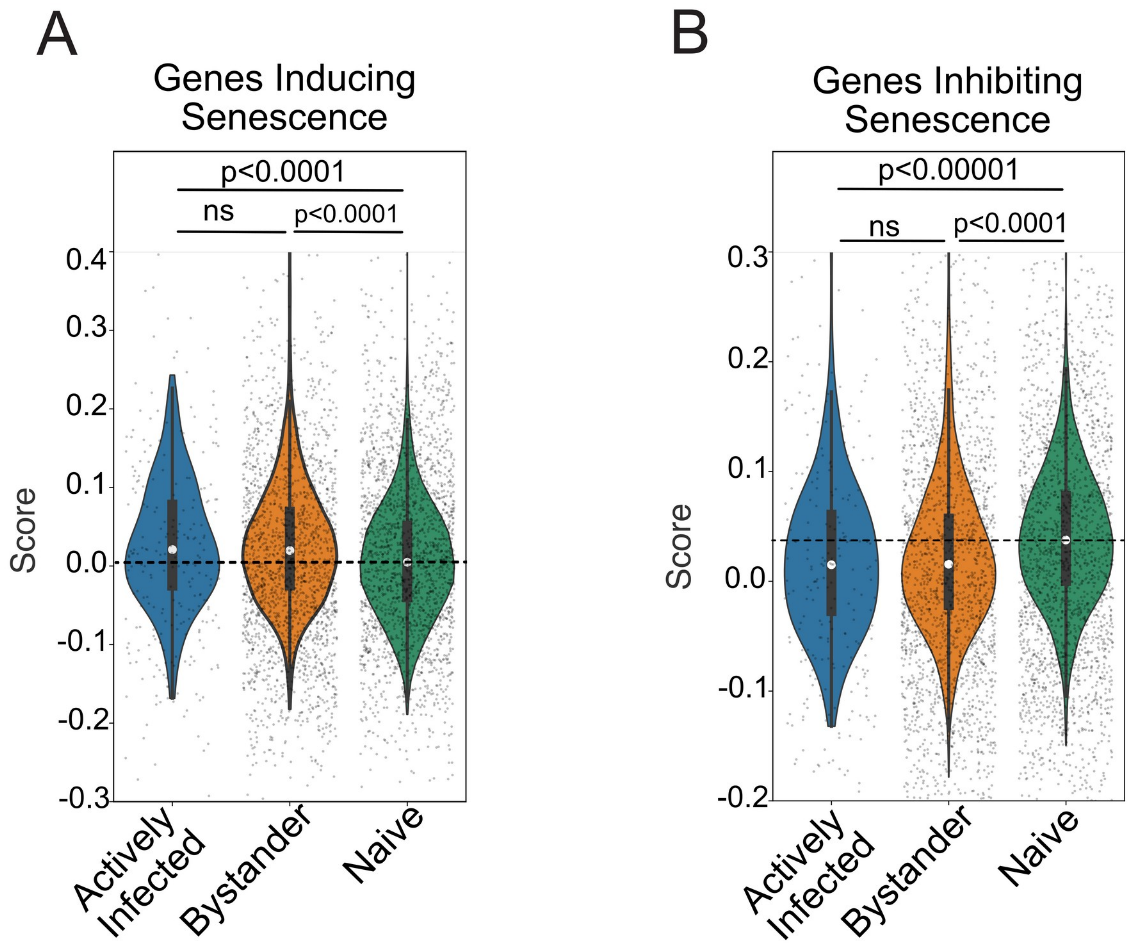

**Supplementary Fig. 22**. Aggregate expression scores of pro- (left panel) and anti-senescence genes (right panel) in naive (saline-treated), bystander (SIVE cells not showing HIV gene expression) and actively infected cells (SIVE cells expressing HIV genes). The entire cell population in the naive, bystander and actively infected groups are used to derive the aggregate scores. The gray dots represent cells. The white dot represent the mean aggregate score for each experimental group.

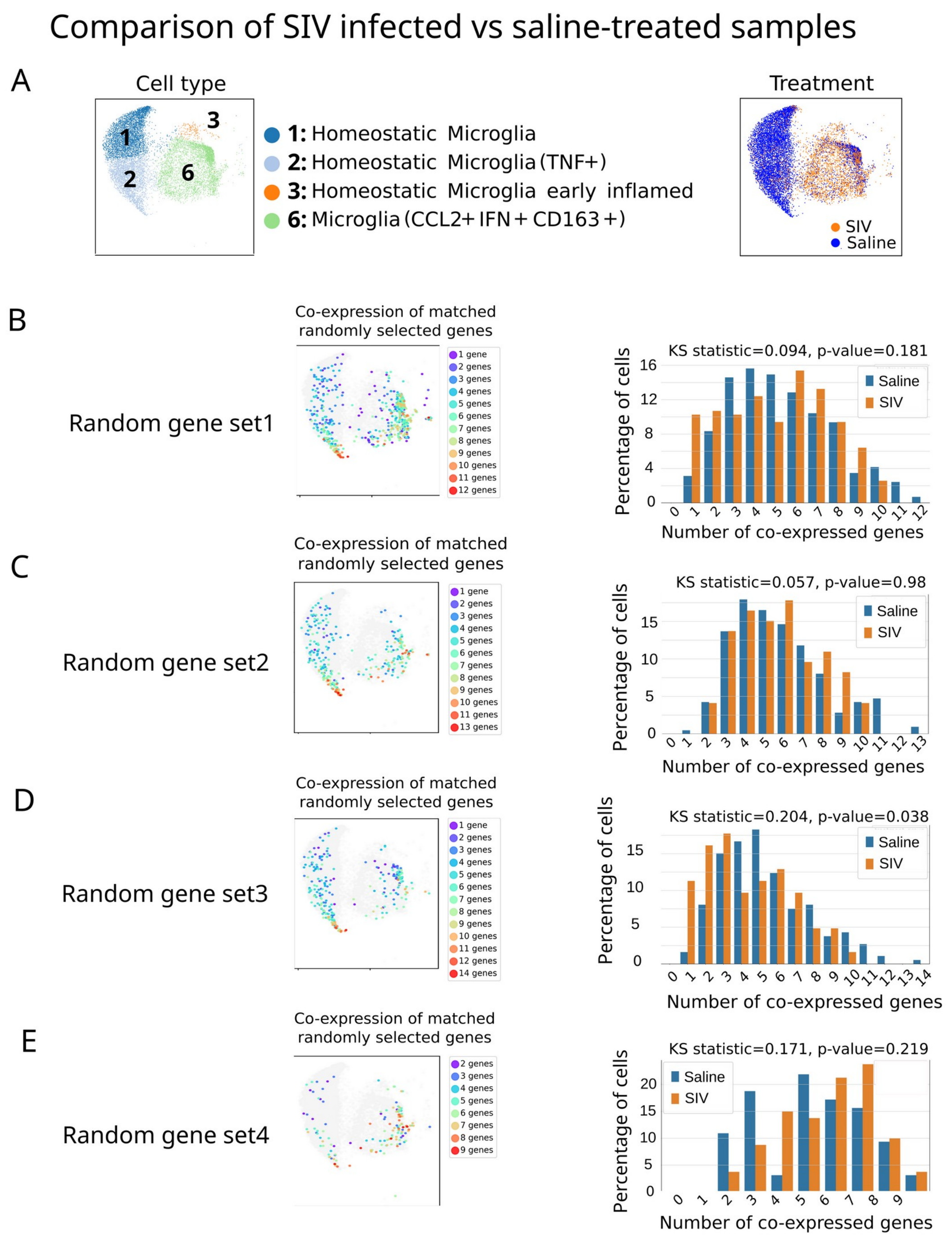

**Supplementary Fig. 23. Control gene sets have a co-expression pattern distinct from that of senescent cells in SIV and saline-treated samples.** A. Distribution of cells across the microglial clusters in the two population under study (SIV and saline-treated). B-E. Pattern of co-expression of four sets of control gene lists (see Methods) are shown, including quantitation of the co-expression patterns on the right and statistical analysis using Kolmogorov-Smirnov test. Dot colors in UMAPs represent the number of co-expressed genes in each cell. For each gene set, bar plots representing the distribution of cells showing different levels of co-expression are shown to the right. Neither of the gene sets showed a statistical significance between SIV and saline groups.

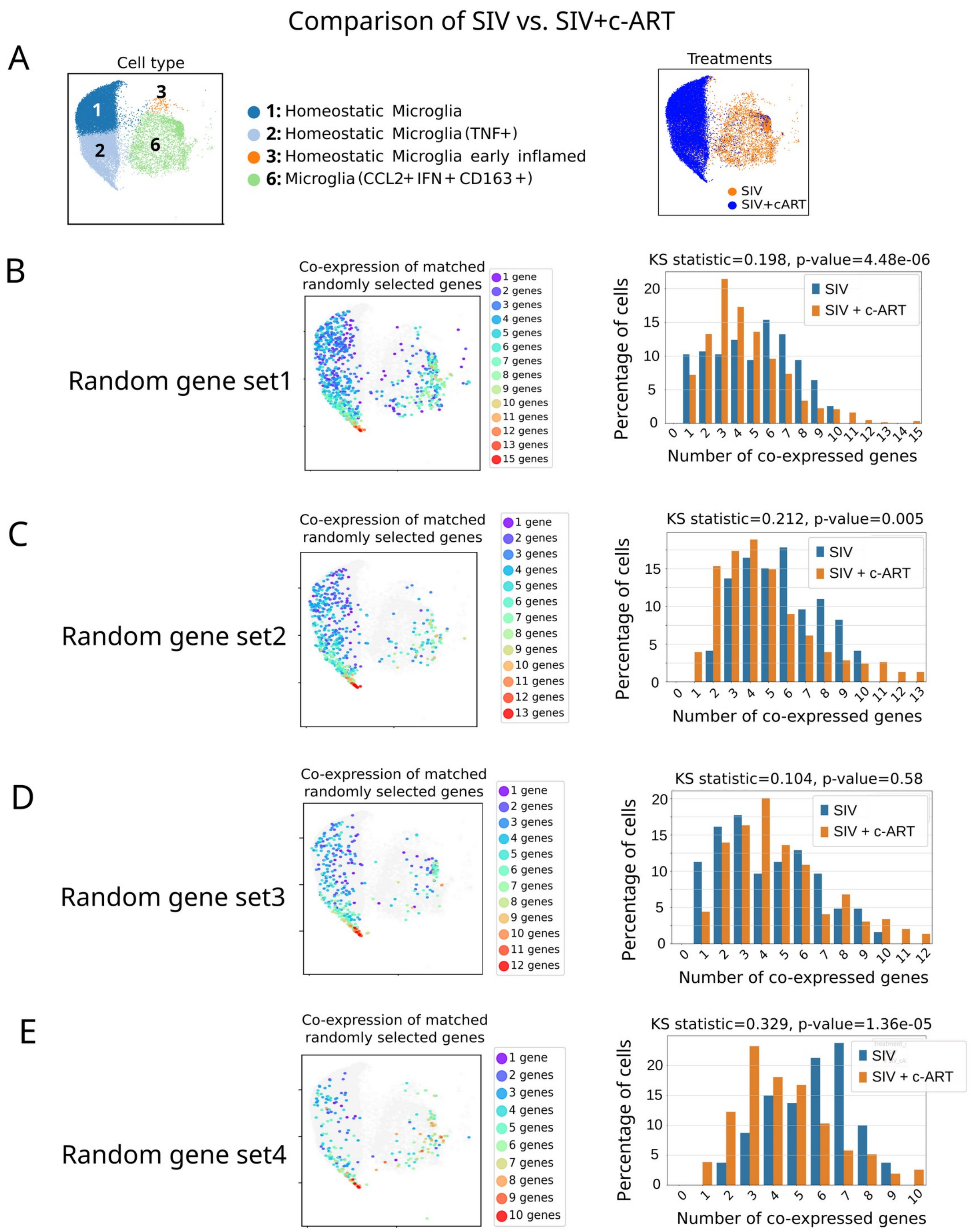

**Supplementary Fig. 24**. SIV-infected cells have a higher number of senescent cells compared to SIV+c-ART samples. A. Distribution of cells across the microglial clusters in the population under study. B-E: The co-expression pattern of four sets of randomly selected cells that match the expression pattern of senescence genes (see Methods). Dot colors in UMAPs represent the number of co-expressed genes in each cell. For each gene set, bar plots representing the distribution of cells showing different levels of co-expression are shown to the right. The results of Kolmogorov–Smirnov test is shown above each bar plot. The relative co-expression pattern of random gene sets in SIV (blue)

and SIV+c-ART (orange) do not resemble the one seen with senescence genes, ruling out the possibility that the observed differences in co-expression of senescence genes is an artifact caused by inherent properties of the dataset and not specific to senescence genes.

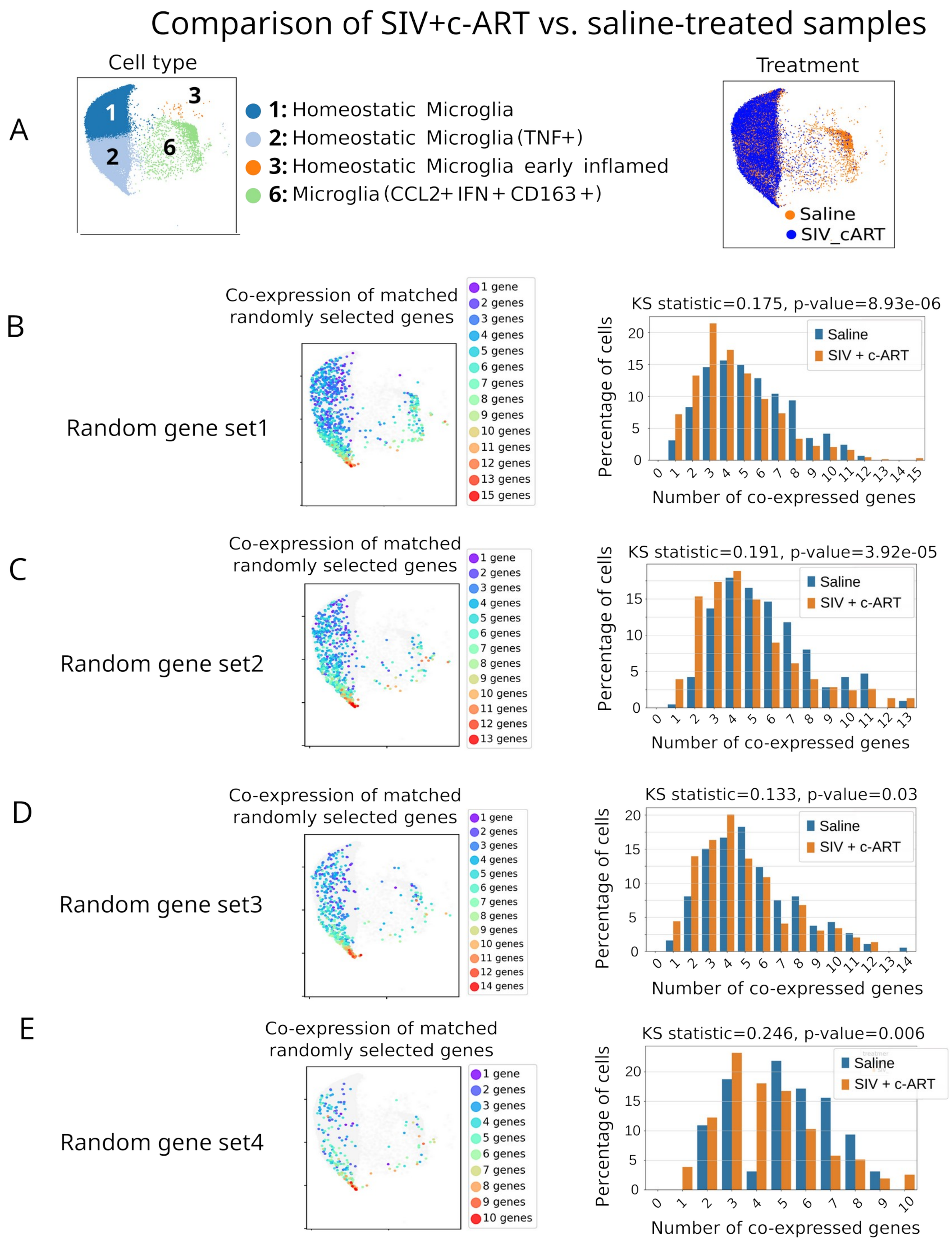

**Supplementary Fig. 25. c-ART treatment following SIV infection leads to reduced inflammation compared to saline-treated samples.** Similar to supplementary figures 23 and 24, four sets of control gene sets are tested for co-expression patterns and quantitated using the barplots on the right for comparison of SIV+cART versus saline samples.

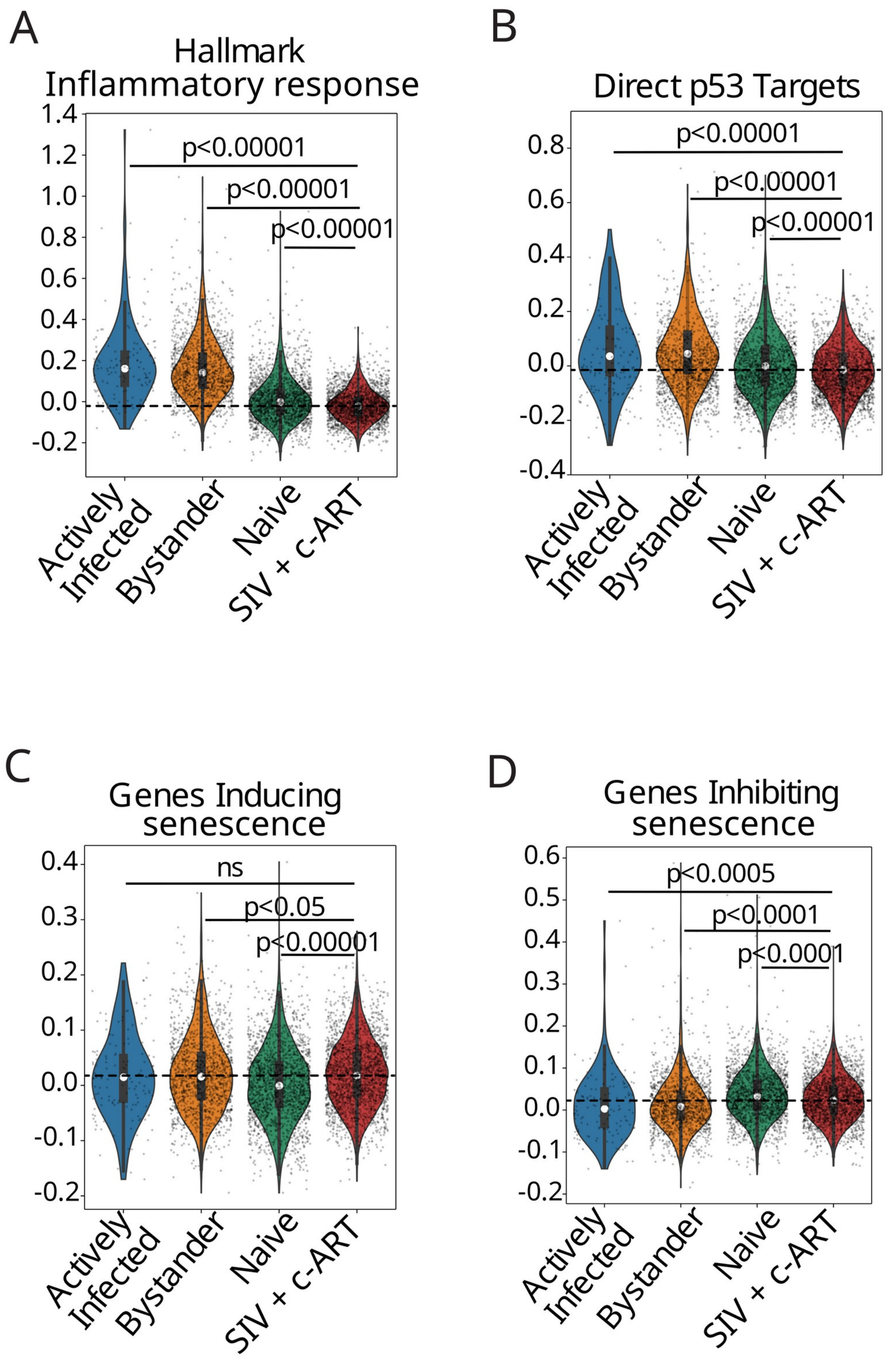

**Supplementary Figure 26. cART does not fully restore SIV infected microglia to homeostatic state.** A and B. Aggregate expression scores for the Hallmark list of inflammatory genes and p53 pathway point to a "hypo-responsive", under-inflamed state in cART-treated microglia relative to homeostatic controls (HIV Naive microglia). C and D. Aggregate expression scores for pro- and anti-senescence genes from the CellAge database points to residual senescence-related changes in cART-treated microglia.

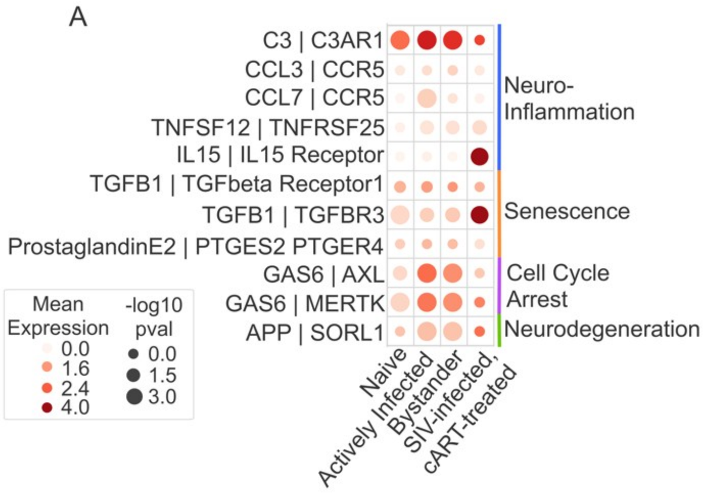

**Supplemental Figure 27.** (A) Dotplots showing the mean expression of outgoing signals from resting/dormant microglia.

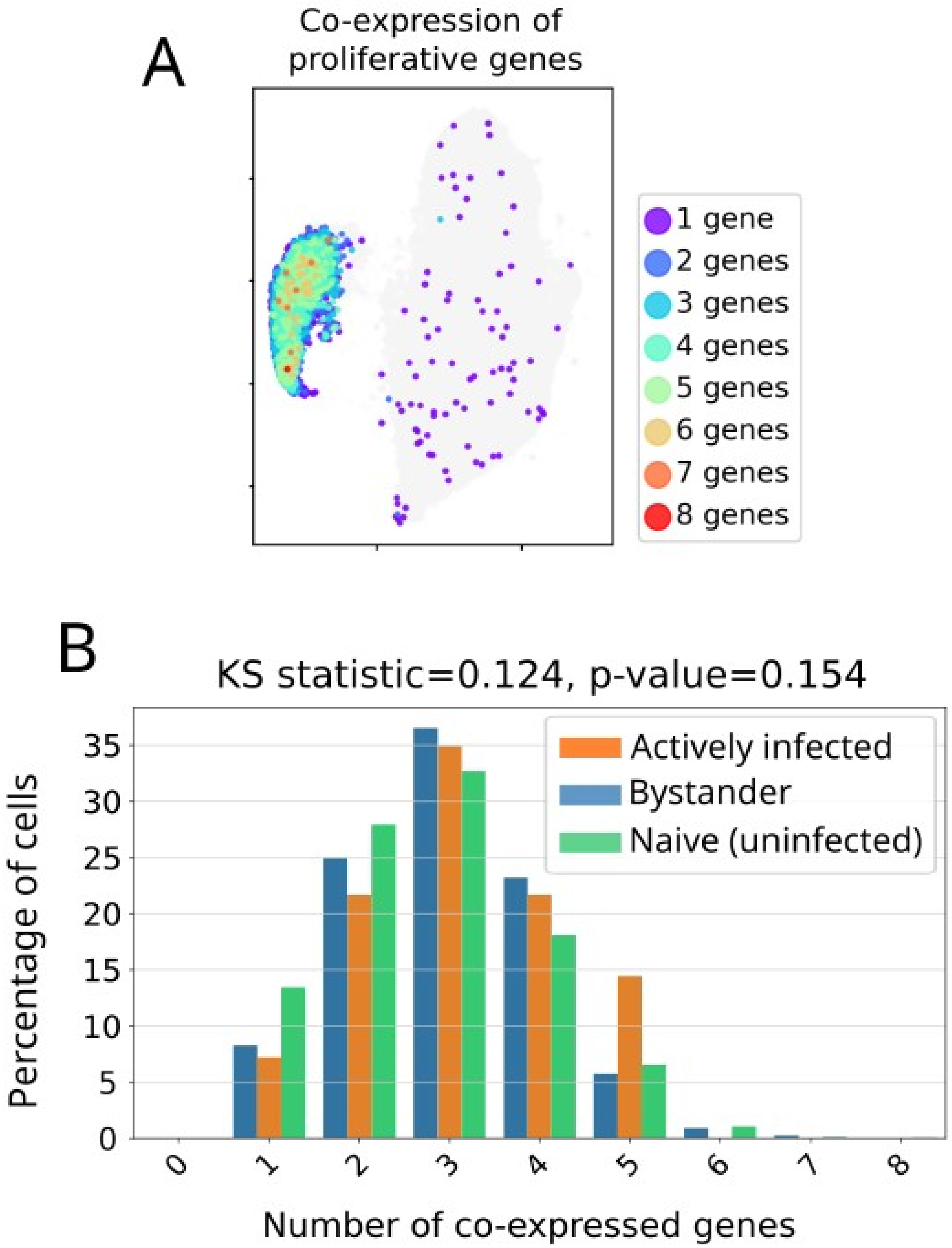

**Supplemental Figure 28. Mild enrichment of expression of proliferative genes in the CDKN2A/p16high cluster in actively infected and bystander cells at day 7 post infection is consistent with a transitional state from post-HIV activated state to senescence.** A. UMAP of co-expression of proliferative genes. B. Distribution analysis of gene co-expression patterns reveals a slight increase in the relative proportion of actively infected and bystander cells that display 3 or more co-expressed genes, although this trend does not achieve statistical significance.
